# Fixation Probabilities of Mutant Alleles in an Ecological Context

**DOI:** 10.64898/2026.07.20.739501

**Authors:** Kunaal Joshi, Susmita Halder, Adrian González Casanova, Michael Lynch

**Author notes:** These authors contributed equally to this work (Kunaal Joshi, Susmita Halder).

## Abstract

For decades, population geneticists have relied upon a formula derived by Malécot and by Kimura to estimate the fixation probability of mutant alleles (*ϕ*). Among other things, this formula leads to the conclusion that in sufficiently large populations *ϕ* asymptotically approaches 2*s*(*N*_*e*_*/N*) (for small *s*), where *s* is the relative selective advantage of the mutant allele, and *N*_*e*_ and *N* respectively denote the effective and absolute (haploid) population sizes. In contrast, in sufficiently small populations, *ϕ* = 1*/N*, with the two domains of behavior being separated at the point where *s ≃* 1*/*(2*N*_*e*_). These results hold when *s, N*, and *N*_*e*_ remain constant during the fixation process, but require modification when populations are changing in size. Here, we show that if there are ecological effects associated with competing alleles, such that the genetic composition of the population influences the total population size, the fixation probability of a beneficial allele can substantially deviate from the levels suggested above, even in the domain of effective neutrality, i.e., |*s*| *≲*1*/*(2*N*_*e*_). We obtain analytical results for a variety of frequency-dependent functions for the overall population size, and show that the overall effect is largely a function of the response at low mutant-allele frequency. We also derive expressions for times to fixation and for fixation probabilities of deleterious alleles, and show how our results relate to prior work for the situation in which population sizes change in a frequency-independent manners.

## Introduction

A key issue in evolutionary genetics is the probability of fixation of a newly arisen mutant allele, as knowledge of this probability combined with the rate of origin of new alleles by mutation provides insight into the potential rate of evolution (Patwa and Wahl, 2008). For decades, population geneticists have relied on a fixation-probability formula developed by Malécot (1952) and Kimura (1957),

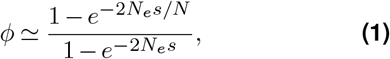

for a haploid population, where *s* is the selective advantage of the mutant relative to the ancestral allele (negative if the mutant is disadvantageous), *N* is the absolute population size (so that the initial frequency of the mutant allele is 1*/N*), and *N*_*e*_ is the effective population size (the effective size determining the rate of genetic drift, which is almost always *< N* owing to various effects of demography, mating systems, and chromosomal architecture; Charlesworth, 2009; Walsh and Lynch, 2018). Note that the formula changes slightly in a diploid population, as a 4 needs to be substituted for the 2 in the denominator of Eq. (1).

Under this model, for situations in which *s ≪* 1*/N*_*e*_, drift overwhelms selection, and the fixation probability is simply equal to the initial frequency, *ϕ ≃*1*/N*. On the other hand, for *s ≫*1*/N*_*e*_, when selection overwhelms drift, the fixation probability is simply equal to twice the selective advantage scaled by the ratio of population sizes, *ϕ ≃* 2*s*(*N*_*e*_*/N*). (In numerous theoretical applications, it is assumed that *N*_*e*_ = *N*, although this is likely almost never the case in biology). Note that the fixation probability is a decreasing function of *N*, so for a given *s*, although increasing *N* is necessary to go from the domain of effective neutrality to effective selection, this increase causes a corresponding decrease in *ϕ*, and selection simply prevents *ϕ* from dropping below 2*s*(*N*_*e*_*/N*) at large *N*_*e*_. This relatively low fixation probability, even with infinite population sizes, occurs because of the very high probability of stochastic loss of any mutant allele in its first few generations, where it is generally present in only a few copies. As *N*_*e*_ is generally smaller than *N* (Charlesworth, 2009; Walsh and Lynch, 2018), conditional on the appearance of a beneficial mutation, 2*s* is typically thought to be the upper limit to the rate of adaptive evolution.

There have been numerous attempts to bring more generality to Equation 1 by introducing additional ecological effects (Slatkin and Hudson, 1991; Griffiths and Tavaré, 1994; Möhle, 2002; Kaj and Krone, 2003; Polanski et al., 2003; Birkner et al., 2009; Steinrücken et al., 2013; Alter and Louzoun, 2016; Terhorst et al., 2016; Koskela and Wilke Berenguer, 2019; Freund, 2020). For example, models have been developed to explore the consequences of populations that are deterministically changing in size (Kimura and Ohta, 1974; Otto and Whitlock, 1997; Waxman, 2011) or fluctuating in size (Ewens, 1967; Otto and Whitlock, 1997; Parsons and Quince, 2007; Ashcroft et al., 2014; Engen et al., 2009; Parsons et al., 2010; Lambert, 2006). The typical assumption in these models is that the change in *N* is governed entirely by external environmental forces, independent of the genetic constitution of the population. Whereas such situations likely exist, the duration of such events may be short relative to the times required for fixation. On the other hand, given that two alleles with selective differences are being evaluated, it seems likely, if not typical, that the two variants will not just differ in survivorship and/or relative fecundity (manifest in *s*) but also in their influences on the environment, which in turn will determine the total numbers of individuals that are supportable.

Here, we consider situations in which alleles alter the environment in ways that equally influence all individuals regardless of their genotypes. This can be expected to happen if alleles depress resources to different extents (independent of their competitive effects with respect to each other, which dictate *s*) or if one of the alleles modifies the environment in other mutually beneficial or deleterious ways. The latter might include “public-goods” sorts of scenarios (e.g., the release of siderophores for nutrient sequestration in microbes, antibiotics or digestive enzymes in fungi, or chemical attractants in invertebrates) or the sharing of signals regarding environmental status (e.g., quorum sensing in bacteria, or warning calls in vertebrates). Such conditioning should also be considered in any attempts to understand the evolutionary genetics of early life, such as pre-cellular settings like an RNA world.

Other distinct genotypic features that influence the physical structure or chemical composition of the environment in ways that are equally experienced can be envisioned here. For example, parasites exist whereby one haplotype modifies the behavior of the host in ways that benefit all occupants and similarly for alternative alleles of a mutualist. Genotypes may also differentially influence the local concentration of an otherwise generally feeding predator, e.g., negatively by the release of chemical deterrents or use of warning coloration, or positively through attraction. In addition, individual size differences associated with allelic effects will result in population-size changes in situations where the environment can only support a fixed amount of biomass. For this reason, a beneficial mutation might even drive a reduction in *N* if organism size increased. A metaphor for the processes envisioned is “a rising tide lifts all boats” (or “a receding tide lowers all boats”). This list is by no means exhaustive.

A particular instance of the model we have in mind involves the case of obligate endosymbionts that influence the metabolic features of their host cells, for instance, mitochondria generating ATP for the host, and thereby influencing the downstream production of cellular building blocks (amino acids, nucleotides, and lipid molecules), and likely modifying cell volume. Because such products will be equally available to all cellular components, a generous endosymbiont genotype would enhance the availability of resources for all other members of the endosymbiont community.

Here, we show that differential (frequency-dependent) influences on resources and other environmental effects (shared by both alleles, and independent of other selective effects) can have significant influences on fixation probabilities. This leads to the conclusion that the potential rate of adaptive evolution can be substantially elevated above the benchmark noted above, although conditions also exist in which fixation is suppressed. Notably, even in relatively small populations experiencing relatively weak selection at the level of survival and fecundity, situations exist in which the fixation probability is elevated above the initial frequency.

## Materials and Methods

### The Models

Here, we consider the conventional situation in which the two allelic types are differentiated by a constant selection coefficient. However, we also allow for the possibility that the alleles interact with their environment in ways that cause shifts in the total population size in response to changes in the genetic constitution of the population. Throughout, we employ a standard Wright-Fisher framework, but unlike the classical model in which *N* is kept constant or changes in a deterministic way, we allow *N* to vary from generation to generation in response to the internal genetic structure of the population, i.e., in a frequency-dependent manner. Depending on the circumstances, the power of drift can then increase or decrease with the advance of a beneficial allele and its influence on the environment. To highlight the focus on the effects of demographic evolution on the fixation probability, we will assume *N*_*e*_ = *N* throughout, though we will also describe a method for applying our results to the case where the effective population size differs from the actual population size.

#### Motivation for population-size formulations

We assume an ancestral haploid population of size *N*_*a*_ and a population size associated with the mutant allele *N*_*m*_ should the latter go to fixation. During a phase of polymorphism, the total population size is assumed to be a function of the frequencies of the two types. This can, in principle, be any arbitrary function *N* (*p*_*m*_) of mutantallele frequency *p*_*m*_ that satisfies the boundary conditions *N* (0) = *N*_*a*_ and *N* (1) = *N*_*m*_. Here, we derive analytical solutions for a general class of models given by,

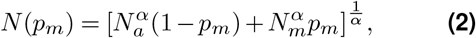

where *α* is an arbitrary parameter (that can take any real value) used to tune the effect of changing mutant-allele frequency on the total population size. We consider four specific functions embodied in Eq. (2), covering a wide range of situations (see Figure 1 and Table 1): the arithmetic mean of the population sizes (AM, corresponding to *α* = 1); the geometric mean (GM, *α→* 0); the harmonic mean (HM, *α* = −1); and the root mean square (RMS, *α* = 2).

**Table 1.**
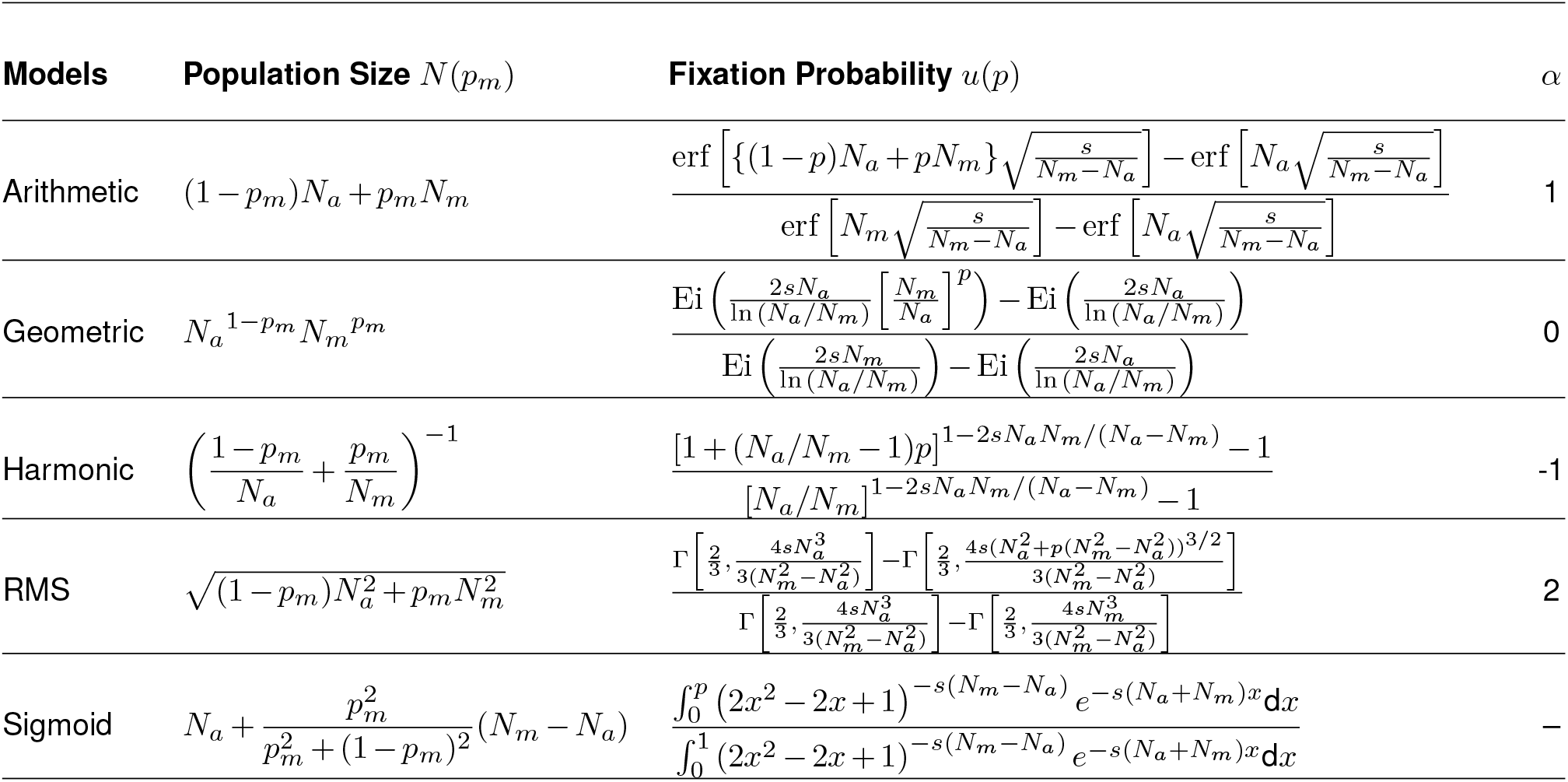
Dependence of the total population size and the probability of fixation on the mutant-allele frequency *p*_*m*_. The Arithmetic, Geometric, Harmonic, and Root Mean Square models are special cases of the general class of models defined in Eq. (2) with tuning parameter *α* values given below. The fixation probability is given as a function of the initial mutant-allele frequency *p*. Here, erf refers to the error function, Ei the exponential integral function, and Γ the upper incomplete Gamma function.

**Figure 1.**
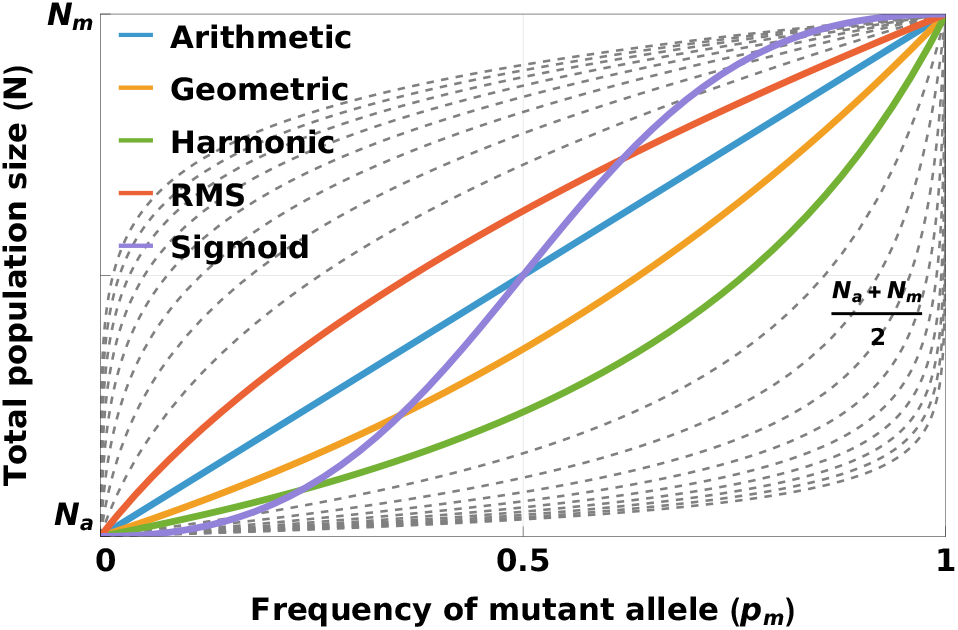
Total population size (*N*) as a function of the mutant frequency (*p*_*m*_) under different models. The general model given by Eq. (2) captures a wide range of behavior, demonstrated here by the dashed lines. Each line represents a different integer value of the tuning parameter *α* in the range −10 to 10. In principle, *α* can take any real value, and as *α* tends to ∞ or −∞, *N* (*p*_*m*_) tends to a step function at 0 or 1 respectively. Specific models with special values of *α* are highlighted: arithmetic mean (*α* = 1), geometric mean (*α* → 0), harmonic mean (*α* = −1), and RMS (*α* = 2) models. In addition, we have considered a sigmoid model that produces a symmetric S-shaped transition. Differences between models are more pronounced when *N*_*a*_ and *N*_*m*_ are far apart.

The arithmetic-mean model assumes that each allele contributes additively to population performance, and (for example) may approximate a public-goods scenario in which each producer contributes additively to the total resource available to the population. The geometric-mean and harmonic-mean models represent situations in which the response to low-frequency alleles is reduced relative to the additive situation, with positive synergistic effects occurring at higher allele frequencies. In contrast, the root-mean-square model represents situations in which population-size increases initially scale rapidly with allele frequency but then exhibit diminishing-returns at high frequencies. In addition, we consider a symmetric sigmoid function, which is not a part of the general class of models tunable with *α*, but combines low-frequency synergistic effects with high-frequency diminishing-return effects. Together, these models cover a wide range of scenarios in which the allele frequency influences the total population size, ranging from relatively weak to relatively strong initial effects. In what follows, we assume a constant coefficient of selection, though in SI Appendix we show that our results can be readily extended to allele-frequency-dependent selection as well.

To derive the analytical results, we used methods inspired by diffusion theory and developed by Kimura (1957) and Kimura and Ohta (1969), which we have detailed in SI Appendix. We later generalize some of the derived results to arbitrary functions beyond those noted above. Note that because the *N* defined by these functions need not be an integer, in the computational implementation of random genetic drift, we applied floor truncation, i.e., chop off the non-integer part of *N* when proceeding to binomial sampling. A related model in which the population size varies due to a difference in the resource consumption efficiencies between the alternate alleles was introduced by González Casanova et al. (2020) and further analyzed by Alsmeyer et al. (2026). focusing on the case where *N*_*m*_ = *O*(*N*_*a*_)). This model gives rise to a population size that is the weighted harmonic mean of the intrinsic population sizes of the allele frequencies, matching our Harmonic Mean model in the diffusion limit (see SI Appendix for details).

### Simulations

In our models, we modify the standard Wright-Fisher framework by allowing the population size *N*, which is kept constant in the standard Wright-Fisher model, to vary from generation to generation as a model-specific function of the mutant-allele frequency. In our simulations, we adhered to a scenario with discrete generations involving consecutive bouts of: 1) selection (governed by the current allele frequency and *s*), converting the old frequency to a new one by multiplying by (1+*s*) divided by the mean population fitness; 2) changing the population size as a function of the mutant-allele frequency after step (1); and 3) drift (governed by the expected frequency following selection and the population size defined by the demographic equations outlined above, and imposing binomial sampling). New mutations initiating as single copies were monitored until reaching frequency 0 (loss) or 1 (fixation). For each set of parameters, data were generated by computer simulations until either 10,000 fixation events occurred (yielding a coefficient of sampling variation of the fixation probability ≃0.01), or until the total number of mutations monitored exceeded 5 *×* 10^10^. As will be shown below, all simulation results are in excellent accord with the derived mathematical expressions.

## Results

### Effect of allele frequency-dependent variation in population size on the probability of fixation

Following prior methods (Kimura and Ohta, 1969), the analytic solution for the fixation probability of the general class of models given by Eq. (2) is

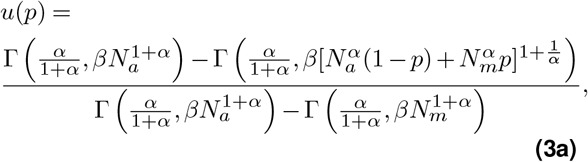

where *p* is the initial frequency of the mutant allele, Γ is the upper incomplete Gamma function, and

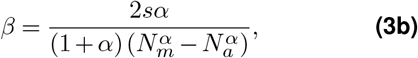

where *s* is the selection coefficient of the mutant allele. This general solution can be further simplified for specific values of *α* (see Table 1, and SI Appendix for derivation).

Simulation results for fixation probabilities as a function of *N*_*a*_ (Figure 2) for a wide range of values of the selection coefficient *s* and relative boost/drop in population size *N*_*m*_*/N*_*a*_ match the derived analytic expressions (Table 1). These analytical results, and those derived in sections that follow, can also be applied to systems in which effective population size differs from actual population size by interpreting *N*_*a*_ and *N*_*m*_ as effective ancestor and mutant population sizes, but retaining the starting frequency *p* in Eq. (3) and Table 1 as the inverse of the actual ancestor population size.

**Figure 2.**
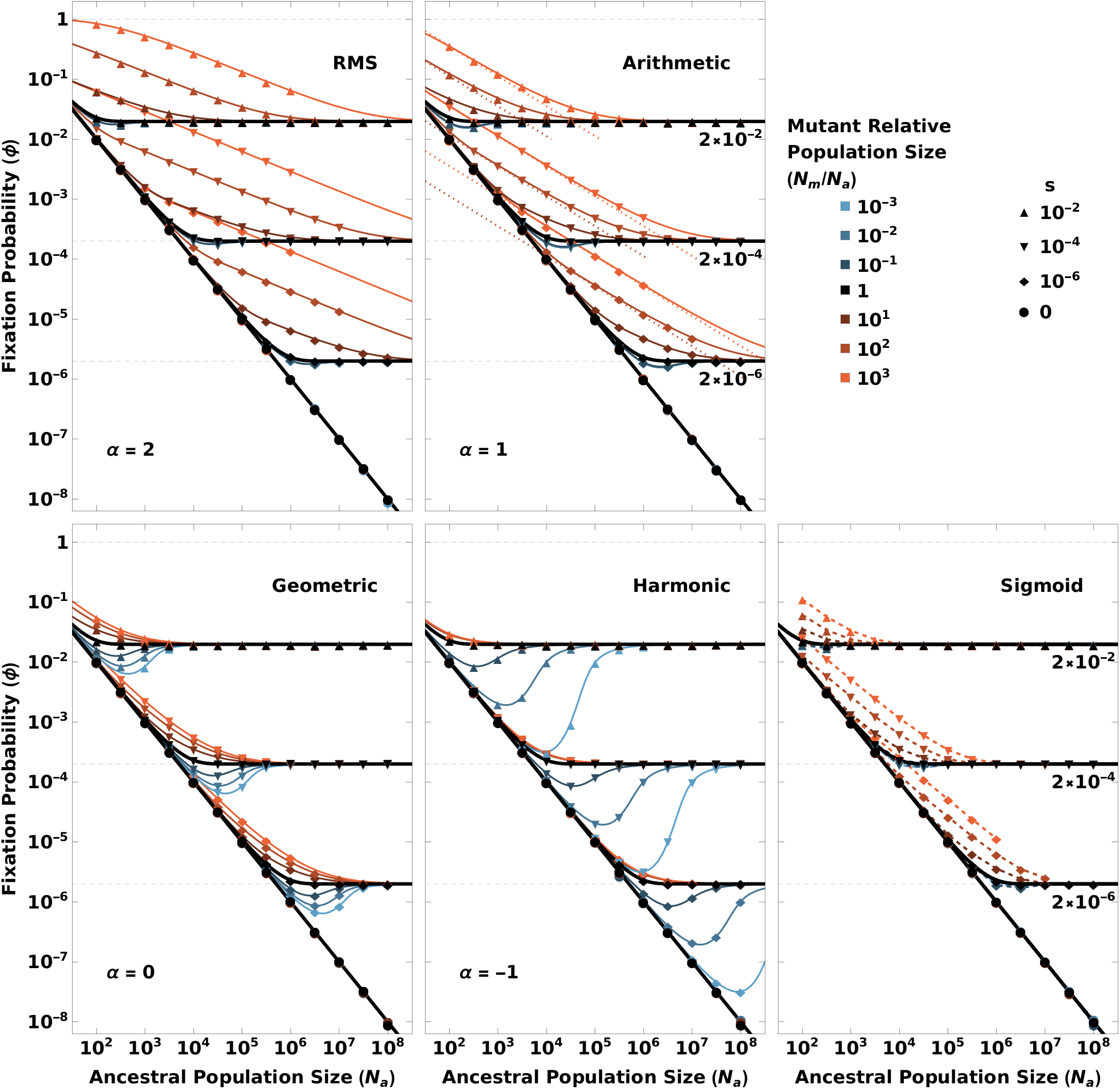
Fixation probabilities of mutant alleles. Fixation probability is plotted as a function of ancestral population size (*N*_*a*_) for different mutant allele-conferred boosts or reductions in population size (*N*_*m*_*/N*_*a*_, shown in different colors) and different selection coefficients *s* (represented by different markers). The solid lines show the analytic expressions from Table 1 where available, while the points (connected by dashed lines for the sigmoid model) represent simulation results. For any given selection coefficient, the fixation probability tends to 1*/N*_*a*_ (solid diagonal line) at small *N*_*a*_ values and approaches 2*s* (dashed horizontal lines) at large *N*_*a*_ values. The black lines in bold show the results for constant population size (*N*_*a*_ = *N*_*m*_). For the arithmetic mean model (second panel), the fixation probability is proportional to 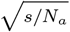 (dotted diagonal lines, with constant of proportionality given by Eq. (4)) at intermediate *N*_*a*_ values (see Table 2). The simulation results are shown for *N*_*m*_ values between 1 and 10^9^.

At the extreme of very small *N* (*N*_*a*_ and *N*_*m*_ *≪*1*/s*), the fixation probability is 1*/N*_*a*_ irrespective of the model, identical to the expectation for the neutral case with constant population size. At the other extreme of very large *N* (*N*_*a*_ and *N*_*m*_ *≫*1*/s*), fixation probabilities all converge to 2*s* irrespective of the model, identical to the strong-selection case with constant population size. However, between these two regimes, the fixation probabilities show strong differences between different models.

**Table 2.**
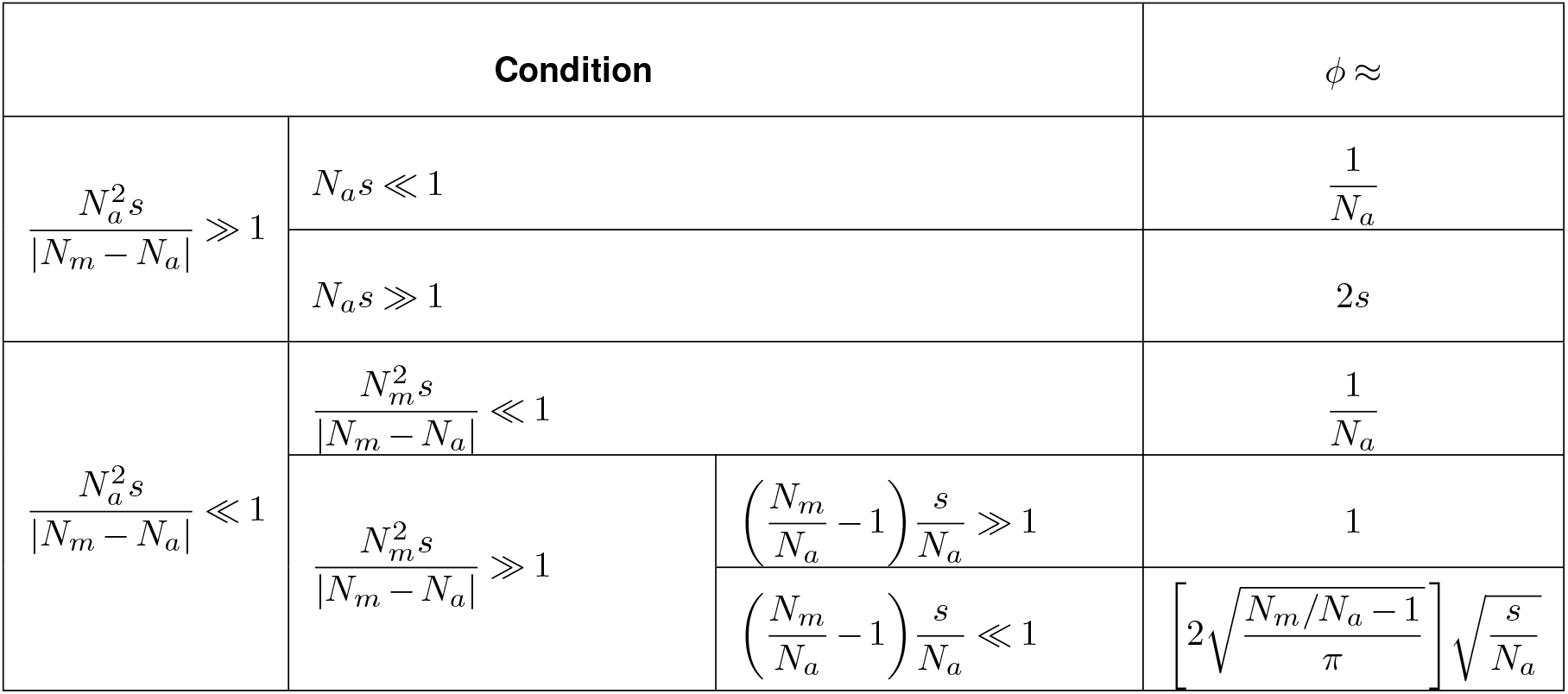
Approximations for the fixation probability *ϕ* for the Arithmetic Mean model under different conditions. The population size as a function of mutant-allele frequency *p*_*m*_ is *N* (*p*_*m*_) = *N*_*a*_ +(*N*_*m*_ − *N*_*a*_)*p*_*m*_ for the Arithmetic Mean model. Here, we consider only beneficial mutations (*s >* 0). Of particular significance is the final condition in the table, under which the fixation probability is boosted to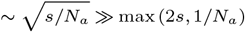, significantly greater than the asymptotic limits for a system with constant population size.

In this intermediate regime, there is asymmetry in the effect of boost versus drop in population size on the fixation probability for different models. For beneficial mutations (positive selection coefficients), for a given *N*_*a*_, the fixation probability is expected to increase with a boost in population size and to decrease with a drop. For positive values of *α*, even a small boost to population size results in a significant increase in fixation probability, but even for significant drops in population size the decrease in fixation probability is negligible. This effect becomes more significant as the magnitude of *α* increases, as demonstrated by comparing the arithmetic mean (*α* = 1) and RMS (*α* = 2) models in Figure 2.

Conversely, for a negative value of *α*, the decrease in fixation probability due to a drop in population size is significant, reaching well below the 2*s* expectation for constant population size, while the increase due to a boost in population size is negligible, as demonstrated by the harmonic mean model (*α* = −1) in Figure 2. The effects of boosts/drops in population size are more balanced for the geometric mean model with *α* = 0. This behavior appears to be a consequence of the property that *α* = 0 forms a critical point at which the log of population size is a linear function of the mutant-allele frequency, and about which it changes behavior from convex (for negative *α* values) to concave (for positive *α* values). The sigmoid model also has a substantial boost effect, but the effect of population size drops is negligible.

The observed differences among arithmetic mean (AM), geometric mean (GM), harmonic mean (HM), and root mean square (RMS) models are consistent with the property that for any set of distinct positive numbers, RMS*>*AM*>*GM*>*HM. Thus, compared to the arithmetic mean, the RMS always lies closer to the upper bound of the set, while the harmonic mean always lies closer to the lower bound, and the geometric mean lies between the harmonic and arithmetic means. Since the population size in any generation is the weighted mean of the ancestral and mutant intrinsic population sizes (*N*_*a*_ and *N*_*m*_), weighted by their respective allele frequencies from the previous generation, consequently, among these models, the effect of a decrease in population size is most pronounced for the harmonic-mean model (the model with the largest *α* in the negative direction), while the effect of an increase is most pronounced for the RMS model (the model with the largest *α* in the positive direction).

Although for both the arithmetic-mean and sigmoid models, the population size as a function of mutant frequency is symmetric (a translated odd function), the arithmetic-mean model has a greater increase in fixation probability due to a more significant early boost in population size. This is a consequence of the slow initial increase in population size for the sigmoid model, which is compensated for by a greater increase when the mutant frequency exceeds 0.5 (see Figure 1). The key point is that the fixation probability is more sensitive to changes in population size when the mutant frequency is close to zero (and hence more vulnerable to stochastic effects) than when the mutant frequency is high.

### Linear approximation for arbitrary dependence of population size on allele frequency

Even for highly beneficial mutations (with large selection coefficient *s*), because frequency at the time of appearance is necessarily close to 0 in large populations, there is a significant chance of being lost by drift in early generations. More specifically, at the time of first appearance, the mutant allele has frequency 1*/N*_*a*_. This causes the population size to change to *≈ N*_*a*_ + *N* ^*’*^ (0)*/N*_*a*_ in the next generation, where *N* (*p*_*m*_) is the function that determines the population size given a mutant frequency *p*_*m*_, and *N* ^*’*^ (0) is the derivative of *N* (*p*_*m*_) at *p*_*m*_ = 0. Thus, the probability of loss in the first generation of sampling is given by

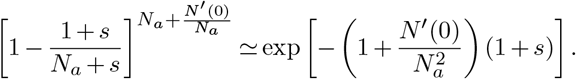

When 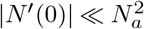, this expression tends to *e*^−(1+*s*)^, which is ≃0.368 when *s ≪*1, imposing an absolute upper bound on the fixation probability of approximately 0.632. In contrast, when 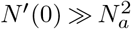, the probability of loss in the first generation becomes negligible, implying that in the limiting case of a very large mutant-conferred population boost during the first generation, the fixation probability could, in principle, approach 1.

If the mutant frequency overcomes this initial hurdle of stochastic loss due to drift at low frequencies, selection is able to pull beneficial mutations to fixation. This is consistent with the previous section’s observation that the initial changes in population size (changes at low mutant frequencies) matter more. Thus, given any arbitrary function for *N* (*p*_*m*_), we propose that its first-order Taylor expansion about 0, *N* (*p*_*m*_) *≈ N* (0) + *N* ^*’*^ (0)*p*_*m*_, is sufficient to determine the fixation probability for small changes in population size.

The analytical solution to this system is identical to the arithmetic mean model (see Table 1), except with relabeled parameters *N*_*a*_ *→N* (0) and *N*_*m*_ *→N* (0) + *N*^*’*^ (0) (since the arithmetic-mean model is also linear, with *N*^*’*^ (0) = *N*_*m*_ − *N*_*a*_). On testing this approximation on our models with non-zero *N*^*’*^ (0), we found it to hold well as long as *N*_*m*_ lies approximately between *N*_*a*_*/*2 and 2*N*_*a*_ (see SI Appendix for details), i.e., as long as the two population sizes are within a factor of two of each other.

### Asymptotic behavior of fixation probability for the linear model

In our general model, for the special case of *α* = 1 (arithmetic mean model), the population size varies linearly with the mutant-allele frequency. Linear models obtained by linearizing arbitrary functions through the Taylor expansion discussed in the previous section are expected to match this arithmetic mean model. Thus, we explore the asymptotic behavior of such models through the arithmetic mean model. For a system with constant population size, the fixation probability tends to 1*/N*_*a*_ when *N*_*a*_ |*s*|*≪* 1 (regime of effective neutrality), and to 2*s* when *N*_*a*_*s ≫*1 (regime of effective selection). For the arithmetic-mean model, the fixation probability tends to these limits under different conditions summarized in Table 2 (see SI Appendix for the derivations).

In addition to the above two regimes, this system exhibits a third regime, when 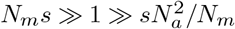 (see Table 2), in which the boost in population size has a significant effect on the fixation probability. An example scenario that falls into this regime is when the ancestral population size is small enough that selective advantage of the mutant allele falls into the regime of effective neutrality (*N*_*a*_*s ≪*1), but the boost in population size brought about by the mutant allele would eventually lead to a crossing over into the regime of effective selection (*N* (*p*_*m*_)*s ≫*1) as the mutant-allele frequency increases. In this regime, the behavior of the fixation probability is further determined by the increase in population size after the appearance of one mutant, *N* (1*/N*_*a*_) − *N* (0) = (*N*_*m*_*/N*_*a*_ − 1). If the quantity (*N*_*m*_*/N*_*a*_ *−* 1)*s/N*_*a*_ *≪* 1, the fixation probability approaches 1, while if this quantity *≪* 1, the fixation probability becomes,

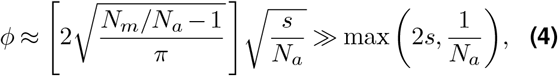

where the inequality follows from the condition 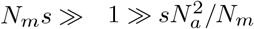. Qualitatively, for a given boost in population size (constant *N*_*m*_*/N*_*a*_), as the ancestral population size *N*_*a*_ increases, the fixation probability changes from 1*/N*_*a*_ at small values of *N*_*a*_, to proportional to 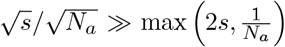 at intermediate values of *N*_*a*_, to 2*s* at large values of *N*_*a*_. The limit in this intermediate regime is shown by the dotted diagonal lines in the second panel of Figure 2. If *N*_*m*_ *≫ N*_*a*_, the probability of fixation in the intermediate regime becomes proportional to 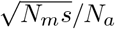. Although the results in Table 2 are presented for the Arithmetic Mean model, they can be extended to any arbitrary function *N* (*p*_*m*_) by linearizing it with Taylor expansion about *p*_*m*_ = 0 (as discussed in the preceding section), and replacing *N*_*a*_ *→ N* (0) and *N*_*m*_ *→ N*^*’*^ (0) + *N* (0).

### Comparison to models in which population size evolves as a function of time

Several prior studies have extended the classical Wright-Fisher model to incorporate a population size that varies as a function of time, independent of mutant-allele frequency (Kimura and Ohta, 1974; Otto and Whitlock, 1997; Ewens, 1967; Ashcroft et al., 2014). One of the earliest of such models is Kimura and Ohta’s logistic growth model (Kimura and Ohta, 1974) in which the population size grows deterministically with time as

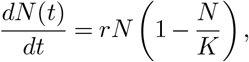

where *K* is the carrying capacity, and *r* is the intrinsic growth rate of the population. We compared the boost in fixation probability under this deterministic time-dependent / frequency-independent model with that when the population size changes purely due to stochastically changing mutant-allele frequency. While our approach and that of Kimura, Ohta, and others are motivated by an interest in how changes in population size affect evolutionary dynamics, our perspective differs in that unlike Kimura and Ohta, we emphasize that population size is not an extrinsic variable but itself subject to evolutionary forces.

The two models are not strongly comparable because the population size at the time of fixation of the mutant allele under Kimura and Ohta’s model is not fixed but depends on the stochastic fixation time. Furthermore, there is no comparable model with a frequency-dependent population size that matches the time dependence of population size in Kimura’s model in the deterministic limit, because the mean rate of change of population size tends to zero when mutant-allele frequency approaches 0 or 1 in models with frequency-dependent population size, while it maintains a constant non-zero value in Kimura’s model (see SI Appendix for details). The frequency-dependent models are thus more significantly influenced by the stochasticity at low mutant-allele frequencies.

We previously noted that it is the initial increase in population size that has the biggest effect on the fixation probability. Thus, for a better comparison, we contrasted Kimura and Ohta’s model with our linear model with the same starting population size *N*_*a*_ and the same initial increase in population size (see SI Appendix for details, and Figure S10 in SI Appendix for results). In the regime of effective selection, the ratio of fixation probabilities of the two models when starting with a single mutant in the population turns out to be (see SI Appendix for details),

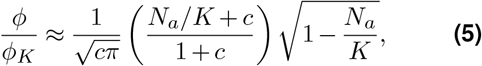

where *ϕ* and *ϕ*_*K*_ are the fixation probabilities for our linear model and Kimura and Ohta’s model respectively, and *c ≡ s/r* is the ratio between the selective advantage of the mutant allele and the intrinsic growth rate of the population under Kimura and Ohta’s model (see Fig. 3 for a visual depiction of when the ratio of fixation probabilities crosses 1 as a function of *c*). From this relation, it can be concluded that when *r ≤ s*, increasing *s* decreases this ratio of fixation probabilities (in this case, the increase in fixation probability in Kimura and Ohta’s model is greater than in our linear model), while increasing *r* increases this ratio (see SI Appendix for details). When *r > s*, there exists a threshold value,

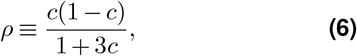

such that if *N*_*a*_*/K < ρ*, then increasing *r* leads to a greater increase in the fixation probability in Kimura and Ohta’s model and increasing *s* leads to a greater increase in our linear model, and vice versa for *N*_*a*_*/K > ρ* (see SI Appendix for details).

**Figure 3.**
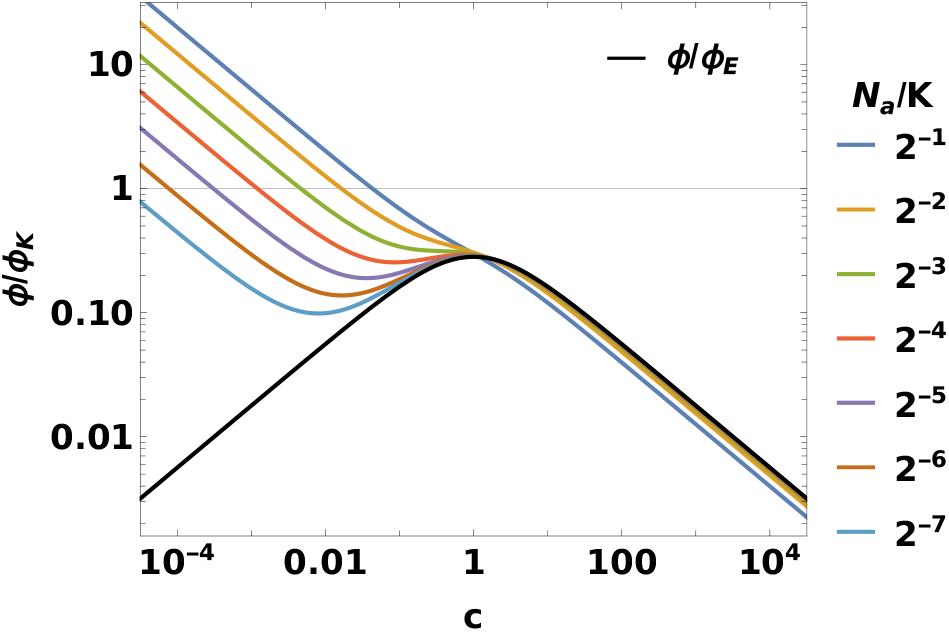
Comparison of fixation probabilities in models with time-dependent versus frequency-dependent population sizes. The ratio of fixation probabilities between our linear model with frequency-dependent population size (*ϕ*) and Kimura’s model with time-dependent population size (*ϕ*_*K*_) is plotted as a function of *c* = *s/r*, the ratio of the selection coefficient to the intrinsic growth rate, for different ratios of initial population size *N*_*a*_ and carrying capacity *K* (see Eq. (5)). Deviations from *ϕ/ϕ*_*K*_ = 1 demonstrate the differences in fixation probabilities between these models. For the fixation probability of the exponential growth model (*ϕ*_*E*_), the ratio *ϕ/ϕ*_*E*_ is plotted in black (see Eq. (7)), and corresponds to *K* → ∞, resulting in *N*_*a*_*/K* → 0.

The results derived here using Kimura and Ohta’s logistic growth model can be extended to the exponential growth model (in which the population size grows exponentially in time independent of the mutant-allele frequency; Otto and Whitlock, 1997) by simply setting *K → ∞*. Thus,

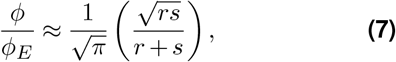

where *ϕ*_*E*_ is the fixation probability for the exponential growth model. The above expression attains a maximum value of 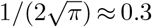 at *r* = *s* (see Fig. 3). Thus, unlike the logistic growth model, which can have a fixation probability greater than or less than our linear model (see Figure S10 in SI Appendix), the fixation probability is always greater for the exponential growth model for the same initial population size and initial increase in population size as our linear model.

We showed above that the fixation probability mainly depends on the rate of population expansion when the mutant allele is at very low frequency, such that models with the same initial population size and initial rate of expansion have similar fixation probabilities. But here we have demonstrated that this similarity is limited to models with population size changing as a function of mutant-allele frequency, and does not extend to situations in which population sizes are changing in time independent of the mutant-allele frequency. The key differences include: (i) The compounding effect that is only present in frequency-dependent models – if the mutant-allele frequency increases, the population size increases in tandem, providing a further boost to advantageous mutations, but conversely, if the mutant-allele frequency falls, the population size also falls, limiting the efficacy of selection. (ii) In addition, in models with a time-dependent population size, the duration for which the mutant allele survives in parallel with the ancestral allele determines the total population sizes available for selection to work on, and the final population size after either fixation or loss of mutation can be very different depending on the absolute time at which fixation or loss occurred, leading to potentially significantly different conditions for recurring identical mutations. In contrast, in frequency-dependent models, the system behavior is independent of external factors such as the absolute time of the arrival of a mutation. (iii) Finally, frequency-dependent models in the deterministic limit can not exhibit the same behavior as the time-dependent models since the mean rate of change of population size tends to zero as the mutant-allele frequency approaches 0 or 1, and are thus more strongly influenced by stochastic effects in these limits.

### Mutants that alter population size without collective benefits

Models with mutant-allele frequency dependent population sizes have been studied in the past primarily in the context of rescue mutations (Wilson et al., 2017; Czuppon et al., 2023). Such systems model situations in which, within an ancestral population experiencing a severe decline in population due to a change in the environmental conditions (such as the introduction of drugs or new competitors, etc.), a mutant arises that is less affected by the current environment and able to maintain higher population numbers in it. The mutants do not provide any collective benefit to the ancestral population, but the total population size (of the ancestors and themselves combined) still increases in an allele-frequency dependent manner, simply because the mutants have a larger carrying capacity. Notable studies have used stochastic birth-death models to find the probabilities of fixation, survival, or establishment (Uecker and Hermisson, 2011; Wilson et al., 2017; Czuppon et al., 2023; Kuosmanen et al., 2025) in such scenarios.

The fixation probability for a special case with time-dependent ancestral per-capita birth and death rates (such that the carrying capacity of the ancestral population decreases exponentially with time), and constant mutant per-capita birth and death rates, was estimated by Uecker and Hermisson (2011) in the application of their more general theory to the rescue mutation problem, using the branching process approach (Fisher, 1923, 1931; Haldane, 1927). This method is limited to the specific cases where the mutation confers a sufficiently large selective advantage such that once it survives the initial period of stochastic fluctuations when at low frequencies, it gets carried to fixation deterministically. Thus, this method limits the analysis to just the initial period where the frequency of a newly arisen mutation is close to 0, and assumes that in this period, the overall population size is solely determined by the ancestral population, without any effects due to the mutants. Thus, such approaches are closer to the time-dependent frequency-independent models discussed in the previous section rather than a true frequency-dependent process.

The limitations of the above approach can be overcome for constant (but different) birth and death rates for both ancestral and mutant subpopulations. Kuosmanen et al. (2025) have shown that in the diffusion limit, irrespective of the relative selective advantage or disadvantage of the mutant allele, such systems converge to the diffusion equation which corresponds to the special case of our general model with *α* = −1 (Harmonic Mean Model) under appropriate parameter transformations (see SI Appendix). In addition, González Casanova et al. (2020) have shown that a system in which the mutant subpopulation has a different efficiency of resource consumption than the ancestral population, thereby leading to a different carrying capacity, also results in the very same diffusion equation. The fixation probability for our *α* = −1 case matches the fixation probabilities derived in these two works under the appropriate parameter transformations.

In the context of drug treatment or immune response to pathogens, to ascertain whether resistant-strain fixing is a cause for concern, it is important to know not only the fixation probability, but also the fixation time, which we analyze in the next section. The fixation time also affects whether the system can be treated as mutation-limited (where the old mutation either fixes or gets wiped out before a new mutation arises), an assumption used by most of the existing works on this subject, such as by Kuosmanen et al. (2025) in their analysis of the longterm evolution of turnover rates for the stochastic birth-death models, among others (Orr and Unckless, 2008; Wilson et al., 2017). In addition, the cases considered above only account for ‘selfish’ mutations, but a more general response could include mutants that benefit the ancestral population too, such as through the production of public goods that counter the drug treatment or suppress the host’s immune response. Such general responses cannot be well described by the models mentioned above, but can be represented by our general model with different *α* values. The responses for specific situations would need to be evaluated experimentally.

### Mean time to fixation

Simulation results for the mean time to fixation as a function of *N*_*a*_ are shown in Figure 4 for different values of the selection coefficient *s* and relative boost/drop in population size *N*_*m*_*/N*_*a*_. For the harmonic mean model, we were able to derive an analytic expression using ideas inspired by diffusion theory. For the remaining models, we were only able to derive analytic expressions for the neutral case (see SI Appendix). Both sets of analytic results (shown in Figure 4) match the simulation results.

**Figure 4.**
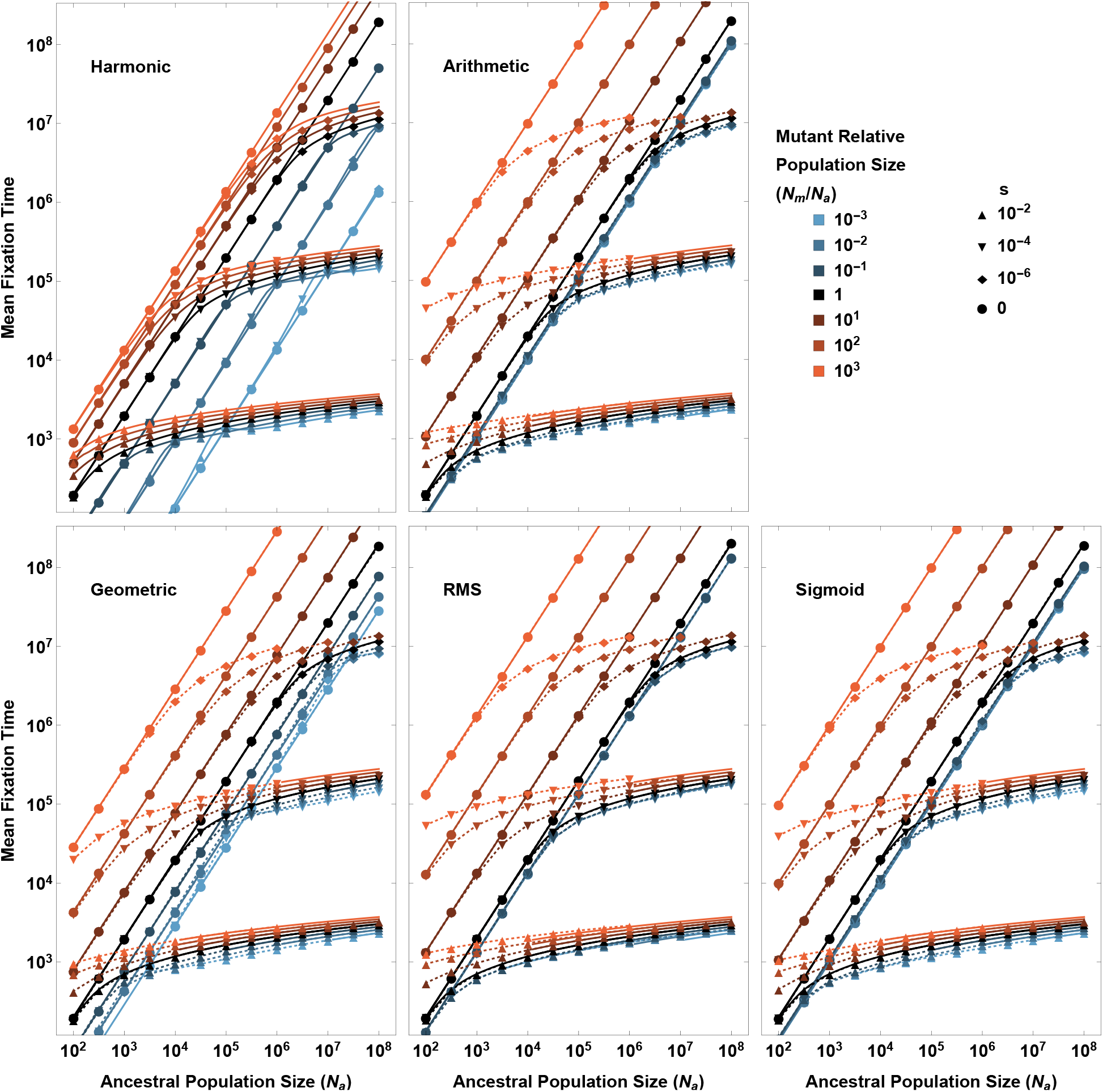
Mean fixation time. Mean fixation time (conditional on fixation) is plotted as a function of ancestral population size (*N*_*a*_) for different mutant allele-conferred boosts or reductions in population size *N*_*m*_*/N*_*a*_ (depicted in different colors) and different selection coefficients *s* (depicted by different markers). The solid lines are the analytic results, while the points (connected by dotted lines) are the simulation results. While the full analytic solution for the harmonic mean model has been obtained, analytic solutions have been derived only for the neutral case (straight diagonal lines) for the other models. In the plots for the Arithmetic, Geometric, RMS, and Sigmoid models, we have also plotted the analytic solution for the harmonic mean model in the regime of strong selection (solid curves, where both *N*_*a*_ and *N*_*m*_ are much greater than 1*/* |*s*|), which matches the simulation results for these models in this regime. The simulation results are shown for *N*_*m*_ values between 1 and 10^9^.

In the regime of effective selection (*N* (*p*_*m*_)*s ≫*1 for all *p*_*m*_), the fixation time becomes model-independent for given ancestral and mutant population sizes. This is because in this regime, the variance in change in mutant frequency per generation (given by *p*_*m*_(1 − *p*_*m*_)*/N* (*p*_*m*_)) becomes negligible compared to the mean change in mutant frequency (given by *sp*_*m*_(1 − *p*_*m*_)). Thus, given that the mutant allele reaches fixation, the mean change in mutant frequency *sp*_*m*_(1 − *p*_*m*_) is what primarily determines the time taken to reach fixation, and it is completely independent of *N* (*p*_*m*_). The population size plays a role only when the term *sp*_*m*_(1 − *p*_*m*_) goes to zero, which happens when *p*_*m*_ *→*0 or *p*_*m*_ *→*1. At these times, the population size is either equal to the ancestral population size *N*_*a*_ (when *p*_*m*_ *→*0) or the mutant population size *N*_*m*_ (when *p*_*m*_ *→*1). Thus, in the strong-selection regime, how the population size varies with mutant frequency is of negligible significance, and only the ancestral and mutant population sizes matter. This is demonstrated in Figure 4, where the simulation results for mean fixation times for the arithmetic, geometric, RMS, and sigmoid models match the analytic expression for the harmonic mean model when both *N*_*a*_ and *N*_*m*_ are ≫1*/* |*s*|.

It has previously been shown that when the population size is constant, mean fixation times starting from any arbitrary mutant frequency are the same for negative and positive selection coefficients of equal magnitude (Maruyama, 1972; Charlesworth, 2020), and we also verify this with our diffusion-theory derived analytic expression for the mean fixation time for constant population size (see SI Appendix). However, this symmetry is altered when the population size varies as a function of mutant-allele frequency. In SI Appendix, we use Kimura and Ohta’s formula (Kimura and Ohta, 1969) to show that for a system with mutant frequency-dependent population size *N* (*p*_*m*_), the mean fixation time for a single, newly arisen mutation with selective advantage *s* is approximately equal to the mean fixation time starting from a single deleterious mutant with selection coefficient − *s* for a system with mutant frequency-dependent population size *N* (1 − *p*_*m*_) (up to a first-order correction term). The two become exactly equal as the starting mutant frequency tends to zero.

## Discussion

The probability of fixation of a mutant allele is one of the most central concepts in evolutionary genetics. In the classical Wright-Fisher model with constant population size, this probability under effective selection (*N*_*e*_*s ≫*1) almost never exceeds a value of 2*sN*_*e*_*/N*, typically a number much smaller than 0.1, and can often be even smaller than this if the effective population size is much less than the absolute size. The traditional models used in population genetics to obtain this result, primarily the Wright-Fisher model (Kimura, 1957; Wright, 1931) and the Moran model (Moran, 1958), generally assume a constant population size. Efforts to extend this work to variable population sizes (Kimura and Ohta, 1974; Otto and Whitlock, 1997; Ewens, 1967; Ashcroft et al., 2014) have focused on the situation in which extrinsic forces drive population-size change deterministically (Kimura and Ohta, 1974; Otto and Whitlock, 1997) or stochastically in time (Otto and Whitlock, 1997; Ewens, 1967; Ashcroft et al., 2014), independent of the genetic constitution of the population. This prior work does show that the fixation probability can exceed the 2*s* benchmark when there is a sustained increase in population size.

We have evaluated a different, albeit not mutually exclusive, scenario in which the total population size is a function of its internal genotypic composition. The effects envisioned here are independent of (and in addition to) the selective differences among potentially competing genotypes. Although one might imagine that selectively advantageous alleles will also lead to larger population sizes, depending on the ecological context, the alternative possibility cannot be ruled out, i.e., in principle, the most selectively advantageous allele might be associated with a reduced population size owing, for example, to increased organism size, higher demands for resources, or modifications of other factors that alter the suitability of the environment for the overall population. The key point is that, when brought to monomorphic states, different genotypes are likely to be associated with overall population-size changes independent of their competitive fitness differences. Our results show that such effects can yield fixation probabilities that are substantially in excess of 2*s*, even exceeding the probabilities for effectively neutral alleles. However, this effect is not universal. Depending on the relationship between population size and allele frequency, situations also exist in which the fixation probability can be reduced below 2*s*, e.g., in the case of harmonic- or geometric-mean dependence.

The extent to which these formal results play out in natural populations, and the degree to which the alternative models in Figure 1 apply, will ultimately need to be examined in empirical settings. So far as we can tell, the assumption of frequency-independent population sizes in prior models has largely been driven by mathematical convenience rather than by direct empirical evidence. However, as noted above, there are innumerable ways in which mutant alleles can have ecological effects that alter overall population sizes. It is for this reason that an exploration of the consequences of frequency-dependent changes in population size seems warranted.

To evaluate the qualitative effects of frequency-dependent population-size change, we developed the analytic solution to a general model with a tuning parameter that determines the degree of response of population size to changing mutant-allele frequencies at different stages of the mutant-allele frequency expansion. We then focused on four specific sub-cases of this general model: arithmetic, geometric, harmonic, and root mean square (RMS), as well as an additional independent sigmoid model. The harmonic approach is related to a model developed in González Casanova et al. (2020) and Alsmeyer et al. (2026), which investigated the situation in which a beneficial mutant decreases population size (*N*_*m*_ *< N*_*a*_) in the regime where the two population sizes are of the same order of magnitude. Together, these models have diverse properties that potentially mimic a variety of ecological scenarios, and different models can yield qualitatively different results. Such differences are primarily a consequence of relative magnitudes of changes in population size during the early stages of expansion of the mutant allele, as the probability of fixation is largely determined by stochastic changes incurred when mutant-allele frequencies are low.

For beneficial mutations that increase population size, a sharper initial increase in population size leads to a greater increase in fixation probability, significantly exceeding the classical threshold of 2*s* in certain cases (such as in the RMS and arithmetic mean models). For the arithmetic mean model, we demonstrate that in situations in which the selectively advantageous mutant allele behaves as effectively neutral at the time of appearance, but over the course of its fixation it increases the population size such that it is no longer effectively neutral, the fixation probability gets boosted to a value proportional to 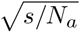 and significantly greater than the classical limits of 2*s* or 1*/N*_*a*_. In contrast, if the initial increase is insignificant, there is hardly any increase in fixation probability (such as in the harmonic mean model), even with the same mutant population size at fixation as other models. Conversely, for beneficial mutations that decrease population size, a sharp initial decrease in population size can cause the fixation probability to fall significantly below the classical threshold (such as in the harmonic mean model), while models with less initial change show hardly any change in fixation probabilities (such as the RMS and arithmetic mean models). These results arise as a consequence of early reductions versus enhancements of the magnitude of random genetic drift.

Despite these differences in fixation probabilities between models, for very large population sizes (when both *N*_*a*_ and *N*_*m*_ are ≫1*/s*), the fixation probability tends to the same 2*s* threshold irrespective of the model. In this regime, although the fixation probability does not depend on the population sizes, the fixation time (conditional on fixation) does. However, because the noise in frequency change is relatively suppressed at intermediate mutant-allele frequencies (relative to the mean change in allele frequencies due to selection), the fixation time is also only weakly influenced by variation in population size at intermediate mutant-allele frequencies. But it is affected by population sizes when the mutant-allele frequency approaches 0 or 1, i.e., when the population size tends to *N*_*a*_ or *N*_*m*_, since the mean change in allele frequencies approaches zero. Thus, in the effective-selection regime, all models have nearly identical fixation times for given values of *N*_*a*_, *N*_*m*_ and *s*.

Our analytical results are largely based on ideas from diffusion theory, which in the mathematical literature might be thought to require that *N*_*m*_ and *N*_*a*_ are of the same order of magnitude. However, for both *N*_*m*_ *≫N*_*a*_ and *N*_*m*_ *≪N*_*a*_, we have found a close correspondence between the predictions of the final formulations and observations from computer simulations. Because this is true for both the fixation probabilities and mean times to fixation of mutant alleles under diverse demographic models, it appears that the diffusion approximations are largely generalizable. Finally, we note that although the main text is primarily focused on the fixation of beneficial mutations under positive selection, our analytic results apply to negative selection as well (see SI Appendix). In addition, we have analyzed a more general version of the arithmetic-mean and RMS models in SI Appendix. We hope that these early results will inspire further mathematical investigation.

The expansion in overall population size due to a new mutation might also be extended to spatial situations in nature, e.g., through the production of public goods that allow the entire population to expand into new regions that previously could not sustain the population. Under such circumstances, the genetic composition and effectiveness of selection at the leading edge of the range expansion might differ due to the phenomenon of gene surfing. This phenomenon has been thoroughly characterized in past studies for populations expanding spatially entirely through external environmental forces, independent of the mutant-allele frequency (Hallatschek and Nelson, 2008; Excoffier and Ray, 2008; Paulose and Hallatschek, 2020; Schlichta et al., 2022). Our work might prove useful in extending such studies to investigate the features of mutant-allele frequency-dependent range expansions.

## ACKNOWLEDGEMENTS

This research was supported by National Institutes of Health award R35-GM122566-01, National Science Foundation award MCB-1518060, Moore and Simons Foundations grant 735927, and Moore Foundation grant 12186.

## AUTHOR CONTRIBUTIONS

ML designed the research. SH, KJ, ML and AGC performed the research. ML performed the simulations. SH and KJ performed the mathematical derivations with assistance from AGC. ML and KJ composed the manuscript with assistance from AGC and SH.

