## Supplementary Information for "Fixation Probabilities of Mutant Alleles in an Ecological Context"

### Supplementary Information for
### The Fixation Probabilities of Mutant Alleles in an Ecological Context

Kunaal Joshi, Susmita Halder, Adrian González Casanova, and Michael Lynch

#### Analytic expressions for the fixation probabilities under the environmental-conditioning model

Here, we attempt to derive some estimators for the fixation probability of a new mutant allele for the situation in which the original and mutant genotypes have different associated population sizes, $N_a$ and $N_m$. The approach is a generalization of Kimura's [1] original derivation (see Appendix 1 of [2] for a review of the approach). The probability of fixation of a mutant allele with initial frequency $p$ is given by,

$$u(p) = \int_0^p G(y)\mathrm{d}y \Big/ \int_0^1 G(y)\mathrm{d}y, \tag{S1}$$

where,

$$G(y) = \exp\left[-\int_0^y \frac{2m(x)}{v(x)}\mathrm{d}x\right],$$

where $m(p_m)$ and $v(p_m)$ are the mean and variance of allele-frequency change over an infinitesimal interval of time. For a haploid population where population size $N(p_m)$ is a function of mutant allele frequency $p_m$, if the mutant allele has a selective advantage $s$, then $m(p_m) = sp_m(1-p_m)$ and $v(p_m) = p_m(1-p_m)/N(p_m)$. We note here that although $p_m$ is a stochastic variable, the population size $N(p_m)$ is a deterministic function of this variable, which allows us to express the mean and variance of allele-frequency change over infinitesimal intervals of time in the above forms. Thus,

$$G(y) = \exp\left[-2s\int_0^y N(x)dx\right]. \tag{S2}$$

For constant population size $N(p_m) = N$, the above reduces to $G(y) = e^{-2Nsy}$, thus $u(p) = \left(1-e^{-2Nsp}\right)/\left(1-e^{-2Ns}\right)$.

##### Frequency dependent selection coefficients

If the selective advantage $s(p_m)$ or disadvantage of the mutant allele over the ancestral allele has an arbitrary dependence on the mutant-allele frequency $p_m$, this dependence can be absorbed into a composite parameter $\eta(p_m)$. From the previous section,

$$G(y) = \exp\left[-2\int_0^y s(p_m)N_e(p_m)dp_m\right] = \exp\left[-2\int_0^y \eta(p_m)dp_m\right],$$

where $N_e$ is the effective population size as a function of mutant-allele frequency, and the composite parameter is given by,

$$\eta(p_m) \equiv s(p_m)N_e(p_m).$$

Thus, our results for fixation probabilities can be extended for frequency dependent selection coefficients by replacing our model's $s$ by 1 and our model's $N(p_m)$ by the actual $s(p_m) * N_e(p_m)$.

##### Arithmetic Mean model

Here,

$$N(p_m) = (1-p_m)N_a + p_m N_m.$$

Replacing this in (S2),

$$G(y) = \exp\left\{-2s\left[\left(y-\frac{y^2}{2}\right)N_a + \frac{y^2}{2}N_m\right]\right\}.$$

This can be integrated by rewriting in quadratic form using,

$$\begin{aligned}\int_0^p G(y)\mathrm{d}y &= \int_0^p \exp\left\{-s[(N_m - N_a)y^2 + 2N_a y]\right\}\mathrm{d}y \\ &= \exp\left\{b^2/a\right\} \cdot \int_0^p \exp\left\{-a[y + (b/a)]^2\right\}\mathrm{d}y,\end{aligned}$$

where $a = (N_m - N_a)s$ and $b = N_a s$. Now, depending on whether $a > 0$ or $a < 0$, the above integral can be written in terms of error functions (erf for $a > 0$ and erfi for $a < 0$),

$$\begin{aligned}\mathrm{erf}(x) =& \frac{2}{\sqrt{\pi}}\int_0^x \exp\left\{-t^2\right\}\mathrm{d}t, \\ \mathrm{erfi}(x) =& \frac{2}{\sqrt{\pi}}\int_0^x \exp\left\{t^2\right\}\mathrm{d}t.\end{aligned}$$

Thus, (S1) results in,

$$u(p) = \begin{cases} \dfrac{\mathrm{erf}\left\{[p + (b/a)]\sqrt{a}\right\} - \mathrm{erf}\left\{(b/a)\sqrt{a}\right\}}{\mathrm{erf}\left\{[1 + (b/a)]\sqrt{a}\right\} - \mathrm{erf}\left\{(b/a)\sqrt{a}\right\}} & \forall\ a > 0, \\ \dfrac{\mathrm{erfi}\left\{[p + (b/a)]\sqrt{-a}\right\} - \mathrm{erfi}\left\{(b/a)\sqrt{-a}\right\}}{\mathrm{erfi}\left\{[1 + (b/a)]\sqrt{-a}\right\} - \mathrm{erfi}\left\{(b/a)\sqrt{-a}\right\}} & \forall\ a < 0. \end{cases}$$

If we allow extension of erf into the complex plane, we can combine the two conditions,

$$u(p) = \frac{\mathrm{erf}\left[\{(1-p)N_a + pN_m\}\sqrt{\frac{s}{N_m - N_a}}\right] - \mathrm{erf}\left[N_a\sqrt{\frac{s}{N_m - N_a}}\right]}{\mathrm{erf}\left[N_m\sqrt{\frac{s}{N_m - N_a}}\right] - \mathrm{erf}\left[N_a\sqrt{\frac{s}{N_m - N_a}}\right]}. \tag{S3}$$

#### Approximations for the arithmetic mean model

We consider the fixation probability for a beneficial mutation ($s > 0$) starting with a single mutant allele ($p = 1/N_a$). Thus, the fixation probability is,

$$\phi = \begin{cases} \dfrac{\mathrm{erf}\left[\left\{N_a + \frac{N_m - N_a}{N_a}\right\}\sqrt{\frac{s}{N_m - N_a}}\right] - \mathrm{erf}\left[N_a\sqrt{\frac{s}{N_m - N_a}}\right]}{\mathrm{erf}\left[N_m\sqrt{\frac{s}{N_m - N_a}}\right] - \mathrm{erf}\left[N_a\sqrt{\frac{s}{N_m - N_a}}\right]} & \forall\ N_m > N_a, \\ \dfrac{\mathrm{erfi}\left[N_a\sqrt{\frac{s}{N_a - N_m}}\right] - \mathrm{erfi}\left[\left\{N_a - \frac{N_a - N_m}{N_a}\right\}\sqrt{\frac{s}{N_a - N_m}}\right]}{\mathrm{erfi}\left[N_a\sqrt{\frac{s}{N_a - N_m}}\right] - \mathrm{erfi}\left[N_m\sqrt{\frac{s}{N_a - N_m}}\right]} & \forall\ N_m < N_a. \end{cases} \tag{S4}$$

The fixation probability can be approximated by different expressions under different regimes:

1. $N_a\sqrt{\left|\frac{s}{N_m - N_a}\right|} \ll 1$ and $N_m\sqrt{\left|\frac{s}{N_m - N_a}\right|} \ll 1$: All erf and erfi terms tend to zero, so we can use the series expansion of $\mathrm{erf}(z)$ and $\mathrm{erfi}(z)$, keeping only the first term, which is $2z/\sqrt{\pi}$ for both. Thus,

$$\phi \approx \frac{1}{N_a}.$$

2. $N_a\sqrt{\left|\frac{s}{N_m - N_a}\right|} \gg 1$ and $N_m\sqrt{\left|\frac{s}{N_m - N_a}\right|} \gg 1$: If $N_m > N_a$, all erf terms tend to 1, so their difference tends to zero. If $N_m < N_a$, all erfi terms tend to $\infty$. In both cases, we can apply L'Hospital's rule and differentiate with respect to $\sqrt{s}$. For $N_m > N_a$,

$$\begin{aligned}\phi \approx& \frac{\left\{N_a + \frac{N_m - N_a}{N_a}\right\}e^{-\left\{N_a + \frac{N_m - N_a}{N_a}\right\}^2\frac{s}{N_m - N_a}} - N_a e^{-\frac{N_a^2 s}{N_m - N_a}}}{N_m e^{-\frac{N_m^2 s}{N_m - N_a}} - N_a e^{-\frac{N_a^2 s}{N_m - N_a}}} \\ =& \frac{1 - \left\{1 + \frac{N_m - N_a}{N_a^2}\right\}e^{-\left\{2 + \frac{N_m - N_a}{N_a^2}\right\}s}}{1 - \frac{N_m}{N_a}e^{-(N_a + N_m)s}} \approx \frac{1 - \left\{1 + \frac{N_m - N_a}{N_a^2}\right\}\{1 - 2s\}}{1 - \frac{N_m}{N_a}e^{-(N_a + N_m)s}} \\ \approx& \frac{2s}{1 - \frac{N_m}{N_a}e^{-(N_a + N_m)s}},\end{aligned}$$

using the property $1 \gg s \gg (N_m - N_a)/N_a^2$. Now, if $N_a s \gg 1$, denominator $\approx 1$, thus $\phi \approx 2s$. But if $N_a s \ll 1$, we get the condition $N_m - N_a \ll N_a^2 s \ll N_a$, thus denominator $\approx 1 - e^{-2N_a s} \approx 2N_a s$, thus $\phi \approx 1/N_a$. Similarly, for $N_m < N_a$,

$$\begin{aligned}\phi \approx & \frac{N_a e^{\frac{N_a^2 s}{N_a - N_m}} - \left\{N_a - \frac{N_a - N_m}{N_a}\right\} e^{\left\{N_a - \frac{N_a - N_m}{N_a}\right\}^2 \frac{s}{N_a - N_m}}}{N_a e^{\frac{N_a^2 s}{N_a - N_m}} - N_m e^{\frac{N_m^2 s}{N_a - N_m}}} \\ = & \frac{1 - \left\{1 - \frac{N_a - N_m}{N_a^2}\right\} e^{-\left\{2 - \frac{N_a - N_m}{N_a^2}\right\} s}}{1 - \frac{N_m}{N_a} e^{-(N_a + N_m)s}} \approx \frac{1 - \left\{1 - \frac{N_a - N_m}{N_a^2}\right\}\{1 - 2s\}}{1 - \frac{N_m}{N_a} e^{-(N_a + N_m)s}} \\ \approx & \frac{2s}{1 - \frac{N_m}{N_a} e^{-(N_a + N_m)s}},\end{aligned}$$

using the property $1 \gg s \gg (N_a - N_m)/N_a^2$. Now, if $N_a s \gg 1$, denominator $\approx 1$, thus $\phi \approx 2s$. But if $N_a s \ll 1$, we get the condition $N_a - N_m \ll N_a^2 s \ll N_a$, thus denominator $\approx 1 - e^{-2N_a s} \approx 2N_a s$, thus $\phi \approx 1/N_a$.

3. $N_a \sqrt{\left|\frac{s}{N_m - N_a}\right|} \ll 1 \ll N_m \sqrt{\left|\frac{s}{N_m - N_a}\right|}$: Here, the denominator becomes $\approx 1$. The value of the numerator depends on whether $\sqrt{\frac{N_m/N_a - 1}{N_a} s} \gg 1$ or $\ll 1$. In the former case, the numerator also becomes $\approx 1$, thus $\phi \approx 1$. In the latter case, the numerator tends to zero, thus we take the first term of the series expansion of erf terms in the numerator, resulting in,

$$\phi \approx 2\sqrt{\frac{(N_m/N_a - 1)s}{\pi N_a}}.$$

4. $N_a \sqrt{\left|\frac{s}{N_m - N_a}\right|} \gg 1 \gg N_m \sqrt{\left|\frac{s}{N_m - N_a}\right|}$: Here,

$$\phi = \frac{1 - \operatorname{erfi}\left[\left\{N_a - \frac{N_a - N_m}{N_a}\right\}\sqrt{\frac{s}{N_a - N_m}}\right] / \operatorname{erfi}\left[N_a \sqrt{\frac{s}{N_a - N_m}}\right]}{1 - \operatorname{erfi}\left[N_m \sqrt{\frac{s}{N_a - N_m}}\right] / \operatorname{erfi}\left[N_a \sqrt{\frac{s}{N_a - N_m}}\right]} \approx 1 - \frac{\operatorname{erfi}\left[\left\{N_a - \frac{N_a - N_m}{N_a}\right\}\sqrt{\frac{s}{N_a - N_m}}\right]}{\operatorname{erfi}\left[N_a \sqrt{\frac{s}{N_a - N_m}}\right]}.$$

Here, both numerator and denominator tend to $\infty$, thus, we apply L'Hospital's rule and differentiate with respect to $\sqrt{s}$. Thus,

$$\phi \approx 1 - \left\{1 - \frac{N_a - N_m}{N_a^2}\right\} e^{-\left\{2 - \frac{N_a - N_m}{N_a^2}\right\} s} \approx 2s,$$

using the property $1 \gg s \gg (N_a - N_m)/N_a^2$.

#### Harmonic-mean model

Here,

$$N(p_m) = \left(\frac{1 - p_m}{N_a} + \frac{p_m}{N_m}\right)^{-1}.$$

Replacing this in (S2),

$$G(p) = \exp\left(\int_0^p -\frac{2sN_a}{1 + ry} \mathrm{d}y\right) = (1 + rp)^{-\frac{2sN_a}{r}},$$

where, $r = (N_a - N_m)/N_m$. Replacing this in (S1),

$$u(p) = \frac{1 - (1 + rp)^{1 - \frac{2sN_a}{r}}}{1 - (1 + r)^{1 - \frac{2sN_a}{r}}}. \tag{S5}$$

#### Geometric Mean model

Here,

$$N(p_m) = {N_a}^{1 - p_m} {N_m}^{p_m}.$$

Replacing this in (S2),

$$G(p) = \exp\left\{-\frac{2sN_a}{\ln\left[N_m/N_a\right]}\left(\frac{N_m}{N_a}\right)^p\right\}.$$

Replacing this in (S1),

$$u(p) = \frac{\mathrm{Ei}\left(-\frac{2sN_m^p N_a^{1-p}}{\ln[\frac{N_m}{N_a}]}\right) - \mathrm{Ei}\left(-\frac{2sN_a}{\ln[\frac{N_m}{N_a}]}\right)}{\mathrm{Ei}\left(-\frac{2sN_m}{\ln[\frac{N_m}{N_a}]}\right) - \mathrm{Ei}\left(-\frac{2sN_a}{\ln[\frac{N_m}{N_a}]}\right)}, \tag{S6}$$

where $\mathrm{Ei}(x)$ is the exponential integral function defined as

$$\mathrm{Ei}(x) = \int_{-\infty}^{x} \frac{e^t}{t} dt$$

for any real non-zero values of x.

#### Sigmoid model

Here,

$$N(p_m) = N_a + \frac{p_m^2}{p_m^2 + (1-p_m)^2}(N_m - N_a).$$

Replacing this in (S2),

$$\begin{aligned} G(p) &= \exp\left(\int_0^p -s\left\{(N_a + N_m) + \frac{(N_a+N_m)(2y-1)}{2y^2-2y+1} - \frac{2(2y-1)N_a}{2y^2-2y+1}\right\} \mathrm{d}y\right) \\ &= \exp\left(-s\left\{(N_a+N_m)p + (N_m - N_a)\ln[2p^2-2p+1]\right\}\right) \\ &= \left(2p^2-2p+1\right)^{-s(N_m-N_a)} \exp\{-s(N_a+N_m)p\}. \end{aligned}$$

Therefore, the fixation probability under the sigmoidal weighting is

$$u(p) \quad = \quad \frac{\int_0^p G(y)\mathrm{d}y}{\int_0^1 G(y)\mathrm{d}y},$$

which needs to be evaluated through numerical integrations.

#### Root Mean Square model

Here,

$$N(p_m) = \sqrt{(1-p_m)N_a^2 + p_m N_m^2}.$$

Instead of the square root, here we derive the fixation probability for the $n$-th power, which ultimately yields the fixation probability for the root mean square model when $n = \frac{1}{2}$ (and arithmetic mean model when $n = 1$).
For the $n$-th power,

$$N(p_m) = \left[(1-p_m)N_a^{\frac{1}{n}} + p_m N_m^{\frac{1}{n}}\right]^n.$$

Thus, $N$ is a concave function of $p_m$ when $0 < n < 1$, and a convex function when $n > 1$. Replacing this in (S2),

$$G(p) = \exp\left(-2s\int_0^p (a_1 + b_1 y)^n \mathrm{d}y\right) = \exp\left\{\frac{2sa_1^{n+1}}{b_1(n+1)}\right\}\exp\left\{-\frac{2s(a_1+b_1p)^{n+1}}{b_1(n+1)}\right\},$$

where, $a_1 = N_a^{\frac{1}{n}},\ b_1 = (N_m^{\frac{1}{n}} - N_a^{\frac{1}{n}})$. Replacing this in (S1),

$$\begin{aligned} u(p) &= \frac{\int_0^p G(y)\mathrm{d}y}{\int_0^1 G(y)\mathrm{d}y} \\ &= 2^{-\frac{1}{n+1}} \frac{a_1(a_1^{n+1}d)^{-\frac{1}{n+1}}\Gamma[\frac{1}{n+1}, 2a_1^{n+1}d] - (a_1+b_1p)(d(a_1+b_1p)^{n+1})^{-\frac{1}{n+1}}\Gamma[\frac{1}{n+1}, 2(a_1+b_1p)^{n+1}d]}{a_1\,\mathrm{E}_{\frac{n}{n+1}}[2da_1^{n+1}] - (a_1+b_1)\,\mathrm{E}_{\frac{n}{n+1}}[2d(a_1+b_1)^{n+1}]} \\ &= \frac{\beta_1^{-\frac{1}{n+1}}\left\{\Gamma[\frac{1}{n+1}, \beta_1 a_1^{n+1}] - \Gamma[\frac{1}{n+1}, \beta_1(a_1+b_1p)^{n+1}]\right\}}{a_1\,\mathrm{E}_{\frac{n}{n+1}}[\beta_1 a_1^{n+1}] - (a_1+b_1)\,\mathrm{E}_{\frac{n}{n+1}}[\beta_1(a_1+b_1)^{n+1}]}, \end{aligned} \tag{S7}$$

where $\beta_1 = 2d = \frac{2s}{b_1(n+1)}$, and $\mathrm{E_n}[x] = \int_1^\infty \frac{e^{-xt}}{t^n} dt$ is the exponential integral function.
Therefore, the fixation probability under the root mean square model is

$$u(p) \quad = \quad \frac{\left(\frac{4s}{3(N_m^2-N_a^2)}\right)^{-\frac{2}{3}} \left\{\Gamma\left[\frac{2}{3}, \frac{4sN_a^3}{3(N_m^2-N_a^2)}\right] - \Gamma\left[\frac{2}{3}, \frac{4s(N_a^2+p(N_m^2-N_a^2))^{\frac{3}{2}}}{3(N_m^2-N_a^2)}\right]\right\}}{N_a^2 \, \mathrm{E}_{\frac{1}{3}}\left[\frac{4sN_a^3}{3(N_m^2-N_a^2)}\right] - N_m^2 \, \mathrm{E}_{\frac{1}{3}}\left[\frac{4sN_m^3}{3(N_m^2-N_a^2)}\right]}. \tag{S8}$$

#### Growth rates

Assuming the population grows exponentially within a generation, here we calculate the growth rates ($R$) required for the population size to change with the change in mutant allele frequency within a generation in accordance with our model.

Within a generation, the mean change in mutant allele frequency is given by $sp_m(1-p_m)$, thus,

$$N(p_m + sp_m(1-p_m)) = N(p_m)e^{R(p_m)}.$$

Plugging in the equation for our general model (Eq. 2),

$$R(p_m) = \frac{1}{\alpha}\log\left(\frac{N_a^\alpha[1-p_m-sp_m(1-p_m)] + N_m^\alpha[p_m+sp_m(1-p_m)]}{N_a^\alpha(1-p_m)+N_m^\alpha p_m}\right) = \frac{1}{\alpha}\log\left(1 + \frac{sp_m(1-p_m)[N_m^\alpha - N_a^\alpha]}{N_a^\alpha(1-p_m)+N_m^\alpha p_m}\right). \tag{S9}$$

Now, to get the bounds on $R$, we use the slope to find local maxima and minima,

$$\begin{aligned}\frac{de^{\alpha R(p_m)}}{dp_m} =& s(N_m^\alpha - N_a^\alpha)\left[\frac{(1-2p_m)[N_a^\alpha(1-p_m)+N_m^\alpha p_m] - (N_m^\alpha - N_a^\alpha)p_m(1-p_m)}{N_a^\alpha(1-p_m)+N_m^\alpha p_m}\right]\\ =& \frac{s(N_m^\alpha - N_a^\alpha)}{N_a^\alpha(1-p_m)+N_m^\alpha p_m}[N_a^\alpha(1-p_m)^2 - N_m^\alpha p_m^2].\end{aligned}$$

This has a zero at,

$$p_m^* = \frac{N_a^{\alpha/2}}{N_a^{\alpha/2}+N_m^{\alpha/2}},$$

with the corresponding growth rate from Eq. (S9),

$$R^* = \frac{1}{\alpha}\log\left(1 + \frac{s(N_m^\alpha - N_a^\alpha)}{\left(N_a^{\alpha/2}+N_m^{\alpha/2}\right)^2}\right).$$

This is the global maximum iff $s(N_m^\alpha - N_a^\alpha) > 0$, i.e., a selectively advantageous mutant increases the population size or a selectively disadvantageous mutant decreases the population size, and the global minimum iff $s(N_m^\alpha - N_a^\alpha) < 0$, i.e., a selectively advantageous mutant decreases the population size, or a selectively disadvantageous mutant increases the population size. But since for the latter case, $R$ is negative, thus the global minimum corresponds to the global maximum of the absolute value of $R$. From Eq. (S9), the other limit on $R$ is 0 when $p_m$ is 0 or 1. Thus, over the course of fixation of the mutant allele, for a deterministic trajectory the growth rate would monotonically change from 0 to $R^*$ then monotonically go back to 0.

#### Convergence of other models to the Harmonic Mean model in the diffusion limit

The diffusion equation for the Harmonic Mean model is given by,

$$dp_m = sp_m(1-p_m)dt + \sqrt{p_m(1-p_m)\left(\frac{1-p_m}{N_a} + \frac{p_m}{N_m}\right)}dW_t,$$

where $W_t$ is the Wiener process. The diffusion equation for the model by Kuosmanen et al. for which they provide the fixation probability is, from Table 1 in [3],

$$dp = \left[s_0 - \left(\frac{\lambda_1^0 - \lambda_2^0}{N}\right)\right]p(1-p)dt + \sqrt{p(1-p)\left[\frac{\lambda_1^0}{N} - p\left(\frac{\lambda_1^0 - \lambda_2^0}{N}\right)\right]}dW_t,$$

where $N$ is constant and $\lambda_i^0$ are the turnover rates of the ancestral ($i = 0$) and mutant ($i = 1$) alleles. This matches our Harmonic Mean model with the following parameter transformations,

$$\begin{aligned} s =& s_0 - \left(\frac{\lambda_1^0 - \lambda_2^0}{N}\right), \\ N_a =& \frac{N}{\lambda_1^0}, \\ N_m =& \frac{N}{\lambda_2^0}. \end{aligned}$$

In model by Gonzalez Casanova et al. [4], the ancestral population and the mutants have different resource consumption efficiencies, which could be independent of their respective selective advantages. With a constant total resource pool available every generation, the total population size in a particular generation is determined by picking ancestral and mutant individuals with probabilities proportional to their respective allele frequencies in the previous generation multiplied by their respective fitnesses until the resource pool is fully consumed. When picked in this manner, the average total population size in a given generation turns out to be the harmonic mean of the ancestral and mutant population sizes weighted by their respective allele frequencies in the previous generation, matching our Harmonic Mean case. The diffusion equation for the model M1 by Gonzalez Casanova et al. is, from Eq. 1.2 in [4],

$$dX_t = -\alpha X_t(1 - X_t)dt + \sqrt{X_t(1 - X_t)(1 - \kappa X_t)}dB_t,$$

where $X_t$ is the mutant allele frequency and $B_t$ is the standard Brownian motion. Scaling time by $N_a$, such that $\tau = N_a t$, we get,

$$dX_\tau = -\frac{\alpha}{N_a} X_\tau(1 - X_\tau)d\tau + \sqrt{X_\tau(1 - X_\tau)\left(\frac{1 - \kappa X_\tau}{N_a}\right)}dB_\tau.$$

This matches our Harmonic Mean model with the following parameter transformations,

$$\begin{aligned} \alpha &= -N_a s, \\ \kappa &= 1 + \frac{N_m}{N_a}. \end{aligned}$$

### Mean Time to Fixation for Harmonic Mean model

Under non-neutrality ($s \neq 0$), obtaining an analytic expression for the mean time to fixation becomes prohibitively difficult. For this case, we focus on deriving an analytical expression for the mean time to fixation only for the harmonic mean case. In contrast, under neutral conditions ($s = 0$), simple analytic expressions can be derived for all models.

From Kimura and Ohta (1969)[5] the mean time to fixation conditioned on fixation starting from frequency $p$ is

$$t(p) = \int_p^1 \Psi(x)u(x)[1 - u(x)]dx + \frac{1 - u(p)}{u(p)} \int_0^p \Psi(x)u^2(x)dx, \tag{S10}$$

where $u(p)$ is the fixation probability given starting frequency $p$, and

$$\Psi(x) = \frac{2\int_0^1 G(x')dx'}{V(x)G(x)}. \tag{S11}$$

For the Harmonic Mean model,

$$G(x) =(1 + rx)^{\alpha-1}, \tag{S12a}$$

$$\int_0^x G(x')dx' =\frac{(1 + rx)^\alpha - 1}{r\alpha}, \tag{S12b}$$

$$V(x) =\frac{(1 + rx)x(1 - x)}{N_a}, \tag{S12c}$$

$$u(x) =\frac{(1 + rx)^\alpha - 1}{(1 + r)^\alpha - 1}, \tag{S12d}$$

where,

$$r \equiv \frac{N_a}{N_m} - 1, \tag{S12e}$$

$$\alpha \equiv 1 - \frac{2N_a s}{r}. \tag{S12f}$$

Thus, $r > -1$ always, and,

$$\Psi(x)u(x) = \frac{2N_a[1-(1+rx)^{-\alpha}]}{r\alpha x(1-x)}. \tag{S13}$$

$$\Psi(x)u(x)[1-u(x)] = A\left[\frac{1-(1+rx)^{\alpha}+(1+r)^{\alpha}[1-(1+rx)^{-\alpha}]}{x(1-x)}\right], \tag{S14}$$

where

$$A = \frac{2N_a}{r\alpha[(1+r)^{\alpha}-1]}. \tag{S15}$$

Also,

$$\Psi(x)u^2(x) = A\left[\frac{(1+rx)^{\alpha}+(1+rx)^{-\alpha}-2}{x(1-x)}\right]. \tag{S16}$$

We solve the required integrals using the following indefinite integrals:

$$\int \frac{dx}{x(1-x)} = \log\left(\frac{x}{1-x}\right), \tag{S17a}$$

$$\begin{aligned}\int \frac{(1+rx)^y}{x(1-x)}dx =& \frac{(1+rx)^{1+y}}{(1+y)(1+r)}\mathrm{F}(1,1+y,2+y,\frac{1+rx}{1+r}) + \frac{1}{y}\left(\frac{1}{rx}\right)^{-y}\mathrm{F}(-y,-y,1-y,-\frac{1}{rx})\\ \equiv& g(y;x).\end{aligned} \tag{S17b}$$

Note that $(1/z)^{-y}$ may not equal $z^y$ when $z$ is negative and $y$ is non-integer. Now, defining

$$h_1(y;x) \equiv \frac{(1+rx)^{1+y}}{(1+y)(1+r)}\mathrm{F}(1,1+y,2+y,\frac{1+rx}{1+r}), \tag{S18a}$$

$$h_2(y;x) \equiv \frac{1}{y}\left(\frac{1}{rx}\right)^{-y}\mathrm{F}(-y,-y,1-y,-\frac{1}{rx}). \tag{S18b}$$

Thus, $g(y;x) = h_1(y;x) + h_2(y;x)$.

$h_1(y;x)$ blows up at $x = 1$ along with $\log(1-x)$, while $h_2(y;x)$ blows up at $x = 0$ along with $\log(x)$. Thus, to integrate Eq.s (S14) and (S16), we have to take the limit after combining terms that blow up. Thus,

$$\begin{aligned}&\lim_{x\to 1^-} -\log(1-x) - h_1(\alpha,x) - (1+r)^{\alpha}[\log(1-x)+h_1(-\alpha,x)]\\ =&[1+(1+r)^{\alpha}]\left[\gamma_E - \log\left(\frac{1}{r}\right) - \log(1+r)\right] + \psi(1-\alpha) + (1+r)^{\alpha}\psi(1+\alpha).\end{aligned} \tag{S19}$$

$$\lim_{x\to 0^+} h_2(\alpha;x) + h_2(-\alpha;x) - 2\log x = 2(\gamma_E + \log r) + \psi(\alpha) + \psi(-\alpha). \tag{S20}$$

Defining

$$\eta(x) \equiv \frac{1-u(x)}{u(x)} = \frac{(1+r)^{\alpha}-(1+rx)^{\alpha}}{(1+rx)^{\alpha}-1}, \tag{S21}$$

$$\begin{aligned}A_0 \equiv& -\frac{1}{(1+\alpha)(1+r)}\mathrm{F}(1,1+\alpha,2+\alpha,\frac{1}{1+r}) - \psi(\alpha) - \psi(-\alpha)\\ &-\frac{1}{(1-\alpha)(1+r)}\mathrm{F}(1,1-\alpha,2-\alpha,\frac{1}{1+r}) - 2\gamma_E + 2\log\left(\frac{1}{r}\right),\end{aligned} \tag{S22}$$

$$\begin{aligned}A_1 \equiv& [1+(1+r)^{\alpha}][\gamma_E + \log r - \log(1+r)] + \psi(1-\alpha) + (1+r)^{\alpha}\psi(1+\alpha)\\ &-\frac{1}{\alpha r^{-\alpha}}\mathrm{F}(-\alpha,-\alpha,1-\alpha,-\frac{1}{r}) + \frac{(1+r)^{\alpha}}{\alpha r^{\alpha}}\mathrm{F}(\alpha,\alpha,1+\alpha,-\frac{1}{r}).\end{aligned} \tag{S23}$$

Now, we can write the solution:

$$t(p) = A\Bigg\{ A_1 + \eta(p)A_0 + [1 + (1+r)^\alpha + 2\eta(p)] \log\left(\frac{1}{p} - 1\right) + [1 + \eta(p)]g(\alpha; p) + [(1+r)^\alpha + \eta(p)]g(-\alpha; p) \Bigg\}. \quad \text{(S24)}$$

### Mean Time to Fixation for Neutral Case

We use the same approach as Kimura and Ohta (1969)[5] to determine the mean fixation time of mutant alleles. Let $u(p,t)$ denote the probability that a mutant allele becomes fixed by the $t$-th generation, given its initial frequency $p$ at time 0. The term $T(p)$ is defined as follows,

$$T(p) = \int_0^\infty t \frac{\partial u(p,t)}{\partial t} dt.$$

This represents the expected total number of generations until the mutant allele becomes fixed, starting from an initial frequency $p$, considering all possibilities, including cases where fixation might not occur. From the above expression, we can derive $t(p)$:

$$t(p) = \frac{T(p)}{u(p)},$$

which represents the average number of generations required for fixation of a mutant allele, given that fixation happens. It is important to note that $u(p)$ stands for the probability of ultimate fixation, expressed as $u(p) = \lim_{t\to\infty} u(p,t)$.

As shown by Kimura (1962), $u(p,t)$ satisfies the following partial differential equation, in which $m(p)$ and $v(p)$ represent the mean and variance of changes in allele frequency, respectively:

$$\frac{\partial u(p,t)}{\partial t} = m(p)\frac{\partial u(p,t)}{\partial p} + \frac{v(p)}{2}\frac{\partial^2 u(p,t)}{\partial p^2}.$$

Differentiating each term of this equation with respect to $t$, multiplying the resulting terms by $t$, and integrating over the interval from 0 to $\infty$, we obtain an ordinary differential equation for $T(p)$ with two boundary conditions: $\lim_{p\to 0} t(p) =$ finite and $t(1) = 0$,

$$T''(p) + a(p)T'(p) + b(p) = 0. \quad \text{(S25)}$$

where $a(p) = \frac{2m(p)}{v(p)}$, $b(p) = \frac{2u(p)}{v(p)}$.
Since $m(p) = 0$ for the neutral case, thus $a(p)$ is also 0, and we only need to consider the $b(p)$ term.

##### For arithmetic case

In this case, $b(p) = \frac{2p\{N_a+(N_m-N_a)p\}}{p(1-p)} = \frac{2\{N_a+(N_m-N_a)p\}}{(1-p)}$.

$$\begin{aligned} T'(p) &= T'(0) + \int_0^p -b(y)dy \\ &= T'(0) - \int_0^p \frac{2\{N_a + (N_m - N_a)y\}}{(1-y)}dy \\ &= T'(0) + 2(N_m - N_a)p + 2N_m \ln[1-p]. \end{aligned}$$

Thus,

$$\begin{aligned} T(p) &= T(1) - \int_p^1 T'(y)dy \\ &= -T'(0)(1-p) - (N_m - N_a)(1-p^2) + 2N_m(p-1)(\ln[1-p] - 1). \end{aligned}$$

Applying the boundary condition $\lim_{p\to 0} T(p) = 0$, we obtain

$$T(p) = (N_a - N_m)(p-1) - (N_m - N_a)(1-p^2) + 2N_m(p-1)\ln[1-p].$$

Therefore, the average time to fixation is

$$t(p) = (N_a - N_m)(1 - \frac{1}{p}) + (N_m - N_a)(p - \frac{1}{p}) - 2N_m(\frac{1}{p} - 1)\ln[1-p]. \quad \text{(S26)}$$

**For harmonic case**

In this case, $b(p) = \frac{2N_a}{(1-p)(1+\frac{N_a-N_m}{N_m}p)} = \frac{2N_a}{(1-p)(1+rp)}$, where $r = \frac{N_a-N_m}{N_m}$.

$$\begin{aligned} T'(p) &= T'(0) - \int_0^p b(y)dy \\ &= T'(0) - \int_0^p \frac{2N_a}{(1-y)(1+ry)}dy \\ &= T'(0) + 2N_a \int_1^{1-p} \frac{1}{z(1+r-rz)}dz \\ &= T'(0) + 2N_m \ln[1-p] - 2N_m \ln[1+rp]. \end{aligned}$$

Thus,

$$\begin{aligned} T(p) &= T(1) - \int_p^1 T'(y)dy \\ &= -\int_p^1 (T'(0) + 2N_m \ln[1-y] - 2N_m \ln[1+ry])dy \\ &= -T'(0)(1-p) + 2N_m(p-1)\ln[1-p]) + \frac{2N_m}{r}\left\{(1+r)\ln[1+r] - (1+rp)\ln[1+rp]\right\}. \end{aligned}$$

Applying the boundary condition $\lim_{p\to 0} T(p) = 0$, we obtain,

$$T(p) = \frac{2N_m}{r}(1+r)p\ln[1+r] + 2N_m(p-1)\ln[1-p] - \frac{2N_m}{r}(1+rp)\ln[1+rp].$$

Therefore, the average time to fixation is

$$t(p) = \frac{2N_m}{r}(1+r)\ln[1+r] - \frac{2N_m}{r}(r+\frac{1}{p})\ln[1+rp] + 2N_m(1-\frac{1}{p})\ln[1-p]. \tag{S27}$$

**For the geometric-mean**

In this case, $b(p) = \frac{2N_a^{1-p}N_m^p}{(1-p)}$,

$$\begin{aligned} T'(p) &= T'(0) + \int_0^p -b(y)dy \\ &= T'(0) - \int_0^p \frac{2N_a^{1-y}N_m^y}{(1-y)}dy \\ &= T'(0) + 2N_m \int_1^{1-p} \frac{(\frac{N_a}{N_m})^x}{x}dx \\ &= T'(0) + 2N_m \left\{\text{Ei}((1-p)\ln[\frac{N_a}{N_m}]) - \text{Ei}(\ln[\frac{N_a}{N_m}])\right\}, \end{aligned}$$

Thus,

$$\begin{aligned} T(p) &= T(1) - \int_p^1 T'(y)dy \\ &= \left\{2N_m \text{Ei}[\ln(\frac{N_a}{N_m})] - T'(0)\right\}(1-p) - 2N_m \int_p^1 \text{Ei}[(1-y)\ln(\frac{N_a}{N_m})]dy. \end{aligned}$$

Applying the boundary condition $\lim_{p\to 0} T(p) = 0$, we obtain

$$T(p) = \frac{-2N_m}{\ln\left(\frac{N_a}{N_m}\right)}\left\{1 - \left(\frac{N_a}{N_m}\right)^{1-p} + (1-p)\ln\left(\frac{N_a}{N_m}\right)\left\{\text{Ei}\left[(1-p)\ln\left(\frac{N_a}{N_m}\right)\right] - \text{Ei}\left[\ln\left(\frac{N_a}{N_m}\right)\right]\right\} + (1-p)\left(\frac{N_a}{N_m}-1\right)\right\}.$$

Therefore the average time to fixation is

$$t(p) = \frac{-2N_m}{p\ln\left(\frac{N_a}{N_m}\right)}\left\{1 - \left(\frac{N_a}{N_m}\right)^{1-p} + (1-p)\ln\left(\frac{N_a}{N_m}\right)\left\{\text{Ei}\left[(1-p)\ln\left(\frac{N_a}{N_m}\right)\right] - \text{Ei}\left[\ln\left(\frac{N_a}{N_m}\right)\right]\right\} + (1-p)\left(\frac{N_a}{N_m}-1\right)\right\}.$$

**For sigmoid case**

In this case,

$$T''(p) = -\frac{2N_a}{1-p} - \frac{2(N_m - N_a)p^2}{(1-p)[p^2 + (1-p)^2]}.$$

Integrating twice,

$$T(p) = -2N_m(1-p)\ln(1-p) + (N_m - N_a)\Big\{(1-p)\Big[\tan^{-1}(1-2p) - \frac{\pi}{4}\Big] - \frac{p}{2}\ln\left[p^2 + (1-p)^2\right]\Big\} + Cp + D,$$

where $C$ and $D$ are integrating constants. Applying the boundary conditions $T(0) = T(1) = 0$, we get $C = D = 0$. Thus, the average time to fixation is

$$t(p) = -2N_m\left(\frac{1}{p} - 1\right)\ln(1-p) + (N_m - N_a)\Big\{\left(\frac{1}{p} - 1\right)\Big[\tan^{-1}(1-2p) - \frac{\pi}{4}\Big] - \frac{1}{2}\ln\left[p^2 + (1-p)^2\right]\Big\}. \tag{S28}$$

**For root mean square**

In this case,

$$T''(p) = -\frac{2\sqrt{(a_1 + b_1 p)}}{1-p},$$

where $a_1 = N_a^2,\ b_1 = (N_m^2 - N_a^2)$.
Instead of deriving the mean fixation time of the square root model, here we attempt to derive the mean fixation time for the model of $n$-th root, which ultimately yields the mean fixation time of the root mean square model when $n = \frac{1}{2}$.

$$T''(p) = -\frac{2(a_1 + b_1 p)^n}{1-p}, \tag{S29}$$

where $a_1 = N_a^{\frac{1}{n}},\ b_1 = (N_m^{\frac{1}{n}} - N_a^{\frac{1}{n}})$.
Integrating twice the above equation and applying boundary conditions $T(0) = T(1) = 0$, we get

$$\begin{aligned} T(p) =& \frac{2p(a_1+b_1)^{2+n}}{b_1(a_1+b_1)(1+n)} - \frac{2(p-1)a_1^{2+n}\Gamma[1+n]_2\tilde{F}_1[1,1+n;3+n;\frac{a_1}{a_1+b_1}]}{b_1(a_1+b_1)} - \frac{2(a_1+b_1p)^{2+n}\Gamma[1+n]_2\tilde{F}_1[1,1+n;3+n;\frac{a_1+b_1p}{a_1+b_1}]}{b_1(a_1+b_1)} \\ =& \frac{2p(a_1+b_1)^{2+n}}{b_1(a_1+b_1)(1+n)} - \frac{2(p-1)a_1^{2+n}\Gamma[1+n]_2F_1[1,1+n;3+n;\frac{a_1}{a_1+b_1}]}{b_1(a_1+b_1)\Gamma[3+n]} - \frac{2(a_1+b_1p)^{2+n}\Gamma[1+n]_2F_1[1,1+n;3+n;\frac{a_1+b_1p}{a_1+b_1}]}{b_1(a_1+b_1)\Gamma[3+n]}. \end{aligned}$$

Thus, the average time to fixation under the nth root model is

$$\begin{aligned} t(p) =& \frac{2(a_1+b_1)^{2+n}}{b_1(a_1+b_1)(1+n)} - \frac{2(1-\frac{1}{p})a_1^{2+n}\Gamma[1+n]_2F_1[1,1+n;3+n;\frac{a_1}{a_1+b_1}]}{b_1(a_1+b_1)\Gamma[3+n]} - \frac{2(a_1+b_1p)^{2+n}\Gamma[1+n]_2F_1[1,1+n;3+n;\frac{a_1+b_1p}{a_1+b_1}]}{pb_1(a_1+b_1)\Gamma[3+n]} \\ =& \frac{2N_m^{\frac{2+n}{n}}}{(N_m^{\frac{1}{n}} - N_a^{\frac{1}{n}})N_m^{\frac{1}{n}}(1+n)} - \frac{2(1-\frac{1}{p})N_a^{\frac{2+n}{n}}\Gamma[1+n]_2F_1[1,1+n;3+n;(\frac{N_a}{N_m})^{\frac{1}{n}}]}{(N_m^{\frac{1}{n}} - N_a^{\frac{1}{n}})N_m^{\frac{1}{n}}\Gamma[3+n]} \\ & - \frac{2\left\{N_a^{\frac{1}{n}} + (N_m^{\frac{1}{n}} - N_a^{\frac{1}{n}})p\right\}^{2+n}\Gamma[1+n]_2F_1[1,1+n;3+n;\frac{\left\{N_a^{\frac{1}{n}} + (N_m^{\frac{1}{n}} - N_a^{\frac{1}{n}})p\right\}}{N_m^{\frac{1}{n}}}]}{p(N_m^{\frac{1}{n}} - N_a^{\frac{1}{n}})N_m^{\frac{1}{n}}\Gamma[3+n]}, \end{aligned}$$

where ${}_2F_1$ is the hypergeometric function.
Therefore, the mean time to fixation of the root mean square model is given by the following after setting $n = \frac{1}{2}$

$$\begin{aligned} t(p) =& \frac{4N_m^5}{3(N_m^2 - N_a^2)N_m^2} - \frac{2(1-\frac{1}{p})N_a^5\Gamma[\frac{3}{2}]_2F_1[1,\frac{3}{2};\frac{7}{2};(\frac{N_a}{N_m})^2]}{(N_m^2 - N_a^2)N_m^2\Gamma[\frac{7}{2}]} \\ & - \frac{2\left\{(N_a^2 + (N_m^2 - N_a^2)p\right\}^{\frac{5}{2}}\Gamma[\frac{3}{2}]_2F_1[1,\frac{3}{2};\frac{7}{2};\frac{\{N_a^2 + (N_m^2 - N_a^2)p\}}{N_m^2}]}{p(N_m^2 - N_a^2)N_m^2\Gamma[\frac{7}{2}]}. \end{aligned} \tag{S30}$$

| Models | Mean Fixation Time $t(p)$ for a neutral mutation |
|---|---|
| Arithmetic | $\{(N_a - N_m)(1-p)\} - \left\{2N_m(\frac{1}{p}-1)\ln[1-p]\right\}$ |
| Geometric | $-\frac{2N_m}{p\ln(N_a/N_m)}\left\{1-\left(\frac{N_a}{N_m}\right)^{1-p} + (1-p)\ln(N_a/N_m)\left\{\mathrm{Ei}[(1-p)\ln(N_a/N_m)] - \mathrm{Ei}[\ln(N_a/N_m)]\right\} + (1-p)\left(\frac{N_a}{N_m}-1\right)\right\}$ |
| Harmonic | $\left\{\frac{2N_m}{r}(1+r)\ln[1+r]\right\} - \left\{\frac{2N_m}{r}(r+\frac{1}{p})\ln[1+rp]\right\} - \left\{2N_m(\frac{1}{p}-1)\ln[1-p]\right\}, \quad \text{where } r = \frac{N_a-N_m}{N_m}$ |
| RMS | $\frac{4N_m^5}{3(N_m^2-N_a^2)N_m^2} - \frac{2(1-\frac{1}{p})N_a^5\Gamma[\frac{3}{2}]_2F_1[1,\frac{3}{2};\frac{7}{2};(\frac{N_a}{N_m})^2]}{(N_m^2-N_a^2)N_m^2\Gamma[\frac{7}{2}]} - \frac{2\left\{(N_a^2+(N_m^2-N_a^2)p\right\}^{\frac{5}{2}}\Gamma[\frac{3}{2}]_2F_1[1,\frac{3}{2};\frac{7}{2};\frac{\left\{(N_a^2+(N_m^2-N_a^2)p\right\}}{N_m^2}]}{p(N_m^2-N_a^2)N_m^2\Gamma[\frac{7}{2}]}$ |
| Sigmoid | $-\left\{2N_m\left(\frac{1}{p}-1\right)\ln(1-p)\right\} + (N_m - N_a)\left\{\left(\frac{1}{p}-1\right)\left[\tan^{-1}(1-2p) - \frac{\pi}{4}\right] - \frac{1}{2}\ln\left[p^2+(1-p)^2\right]\right\}$ |

Table S1: Mean Time to Fixation for neutral case

### Relation between mean fixation times with positive and negative selection

From Kimura and Ohta, 1969 [5], the mean time to fixation starting from frequency $p$ is,

$$t(p) = \int_p^1 \Psi(x)u(x)[1-u(x)]dx + \frac{1-u(p)}{u(p)}\int_0^p \Psi(x)u^2(x)dx, \tag{S31}$$

where $u(p)$ is the fixation probability given starting frequency $p$, and

$$\Psi(x) = \frac{2\int_0^1 G(x')dx'}{V(x)G(x)}, \tag{S32a}$$

$$u(x) = \frac{\int_0^x G(x')dx'}{\int_0^1 G(x')dx'}, \tag{S32b}$$

$$G(x) = e^{-2\int \frac{M_{\delta x}}{V_{\delta x}}dx}. \tag{S32c}$$

When population size varies with mutant frequency $x$ as $N = f(x)$, we have $M_{\delta x} = sx(1-x)$ and $V_{\delta x} = x(1-x)/f(x)$. Defining

$$F(x) \equiv \int_0^x f(x)dx, \tag{S33}$$

thus,

$$G(x) = e^{-2sF(x)}, \tag{S34a}$$

$$u(x) = \frac{\int_0^x e^{-2sF(x_1)}dx_1}{\int_0^1 e^{-2sF(x_1)}dx_1}, \tag{S34b}$$

$$\Psi(x)u(x) = \frac{2f(x)\int_0^x e^{-2sF(x_1)}dx_1}{x(1-x)e^{-2sF(x)}}. \tag{S34c}$$

Thus,

$$t(p) = \int_p^1 \frac{2f(x)\left[\int_0^x e^{-2sF(x_1)}dx_1\right]\left[\int_x^1 e^{-2sF(x_1)}dx_1\right]}{x(1-x)e^{-2sF(x)}\int_0^1 e^{-2sF(x_1)}dx_1}dx \\ + \frac{\int_p^1 e^{-2sF(x_1)}dx_1}{\int_0^p e^{-2sF(x_1)}dx_1}\int_0^p \frac{2f(x)\left[\int_0^x e^{-2sF(x_1)}dx_1\right]^2}{x(1-x)e^{-2sF(x)}\int_0^1 e^{-2sF(x_1)}dx_1}dx. \tag{S35}$$

$$u'(x) = \frac{e^{-2sF(x)}}{\int_0^1 e^{-2sF(x_1)}dx_1} \tag{S36}$$

Now, as $u(0) = 0$, using L'Hospital's rule,.

$$\lim_{p\to 0}\frac{\int_0^p \Psi(x)u^2(x)dx}{u(p)} = \lim_{p\to 0}\frac{2f(0)\left[\int_0^p e^{-2sF(x)}dx\right]^2}{p} = 0. \tag{S37}$$

Thus,

$$t(0) = \int_0^1 \frac{2f(x)\left[\int_0^x e^{-2sF(x_1)}dx_1\right]\left[\int_x^1 e^{-2sF(x_1)}dx_1\right]}{x(1-x)e^{-2sF(x)}\int_0^1 e^{-2sF(x_1)}dx_1}dx. \tag{S38}$$

Now,

$$\begin{aligned} t'(0) &= \lim_{p\to 0} -\Psi(p)u(p) - \frac{u'(0)}{u^2(p)}\int_0^p \Psi(x)u^2(x)dx + \Psi(p)u(p) \\ &= \lim_{p\to 0} -\frac{\Psi(p)u(p)}{2} = \lim_{p\to 0} -\frac{f(0)\int_0^p e^{-2sF(x)}dx}{p} \\ &= -f(0). \end{aligned} \tag{S39}$$

Thus, if population size is large and we start from one mutant in population,

$$t\left(\frac{1}{f(0)}\right) \approx t(0) + t'(0)\frac{1}{f(0)} = t(0) - 1. \tag{S40}$$

Now, defining

$$h(x) \equiv f(1-x), \tag{S41a}$$

$$H(x) \equiv \int_0^x h(x_1)dx_1. \tag{S41b}$$

Thus,

$$\begin{aligned} F(1-x) &= \int_0^{1-x} f(x_1)dx_1 = \int_1^{1-x} f(x_1)dx_1 + \int_0^1 f(x_1)dx_1 \\ &= -\int_0^x h(y_1)dy_1 - \int_1^0 h(y_1)dy_1 = H(1) - H(x) \end{aligned} \tag{S42}$$

$$\begin{aligned} \int_0^{1-x} e^{-2sF(x_1)}dx_1 &= \int_x^1 e^{-2sF(1-y_1)}dy_1 \\ &= e^{-2sH(1)}\int_x^1 e^{2sH(y_1)}dy_1. \end{aligned} \tag{S43}$$

$$\begin{aligned} \int_{1-x}^1 e^{-2sF(x_1)}dx_1 &= \int_0^x e^{-2sF(1-y_1)}dy_1 \\ &= e^{-2sH(1)}\int_0^x e^{2sH(y_1)}dy_1. \end{aligned} \tag{S44}$$

Thus, replacing $y = 1 - x$ in Eq. (S38),

$$t(0) = \int_0^1 \frac{2h(y)\left[e^{-2sH(1)}\int_y^1 e^{2sH(y_1)}dy_1\right]\left[e^{-2sH(1)}\int_0^y e^{2sH(y_1)}dy_1\right]}{y(1-y)e^{-2s[H(1)-H(y)]}e^{-2sH(1)}\int_0^1 e^{2sH(y_1)}dy_1}dy$$

$$= \int_0^1 \frac{2h(x)\left[\int_0^x e^{2sH(x_1)}dx_1\right]\left[\int_x^1 e^{2sH(x_1)}dx_1\right]}{x(1-x)e^{2sH(x)}\int_0^1 e^{2sH(x_1)}dx_1}dx \tag{S45}$$

This is identical to Eq. (S38), except with sign of $s$ changed. Thus, mean fixation time starting from one mutant for a system with mutant frequency dependent population size $f(x)$ and selective advantage of mutant $s$ is approximately equal to mean fixation time starting from one mutant for a system with mutant frequency dependent population size $f(1-x)$ and selective advantage of mutant $-s$ (equal upto first order correction term). The two become exactly equal as starting mutant frequency goes to zero.

### Testing the linear approximation on models with non-zero initial slope

For our different models with different $N(p_m)$, we compare the analytic expression of fixation probabilities derived above with the analytic expression derived from the linear approximation $N(p_m) \approx N(0) + N'(0)p_m$, where $N'(p_m)$ is the derivative of the population size as a function of mutant allele frequency. $N'(0) = 0$ for the sigmoid model, thus we exclude it from comparison. We also exclude the arithmetic mean model, because for this model this approximation is exact, thus there is no need for comparison. For the geometric mean and RMS models, the results match closely in the range $N_a/2 \leq N_m \leq 2N_a$ (see Figures S1 and S2). For the harmonic mean model, $N(0) + N'(0) = 0$ when $N_m = 2^{-1}N_a$, which results in an indeterminate result, thus we limit our range to $2^{-0.8}N_a \leq N_m \leq 2N_a$ (see Figure S3).

### Testing the analytic expressions of fixation probability and mean fixation time for negative values of selection coefficient

We have compared the analytic expressions for the fixation probability for the geometric mean model, harmonic mean model, and our general $n$-th root model (with $n = 1$, corresponding to the arithmetic mean case) with simulation results with negative selection coefficients in Figure S4. We have also compared the analytic expression for mean fixation time for the harmonic mean model with simulation results with negative selection coefficients in Figures S5 and S6. The analytic expressions precisely match the simulation results.

### Testing the relation between mean fixation times with positive and negative selection

We have matched simulation results for the mean fixation time for the harmonic mean, arithmetic mean, geometric mean, and sigmoid models with positive selection coefficients $s$, with the simulation results for these corresponding models with negative selection coefficients of the same magnitude after interchanging $N_a$ and $N_m$ (see Figures S6, S7, S8, S9). These results precisely overlap.

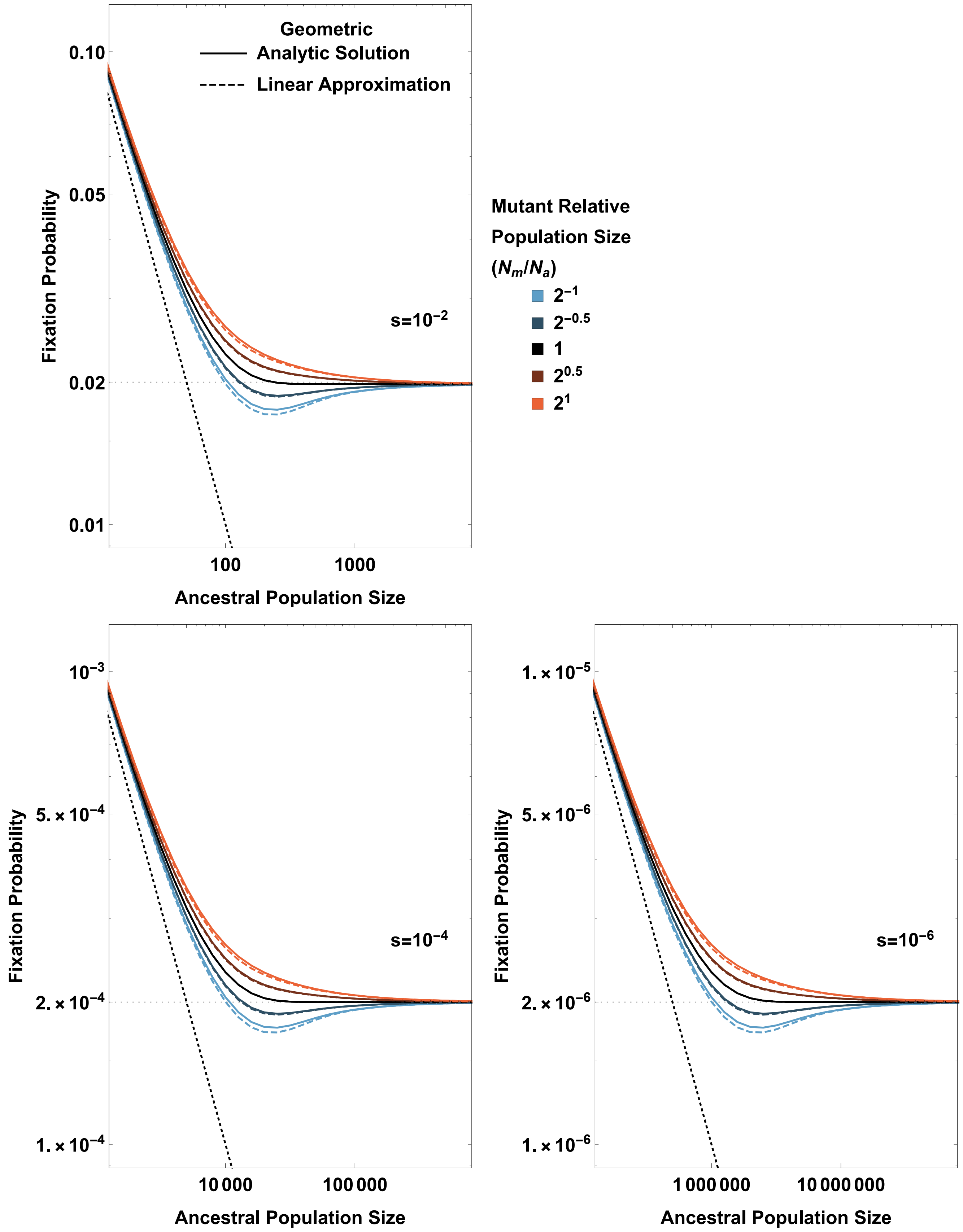


Figure S1: Fixation probability ($\phi$) as a function of ancestral population size ($N_a$) for different values of the selection coefficients $s$ and relative boost (or drop) in population size ($N_m/N_a$), for the geometric mean model. The solid lines represent the analytical results, while the dashed lines represent the linear approximation results; both are closely matched with each other. The gray dotted line represents $\phi = 2s$ (the asymptotic limit of the fixation probability under positive selection at large population sizes), and the black dotted line represents $\phi = 1/N$ (the fixation probability under neutrality).

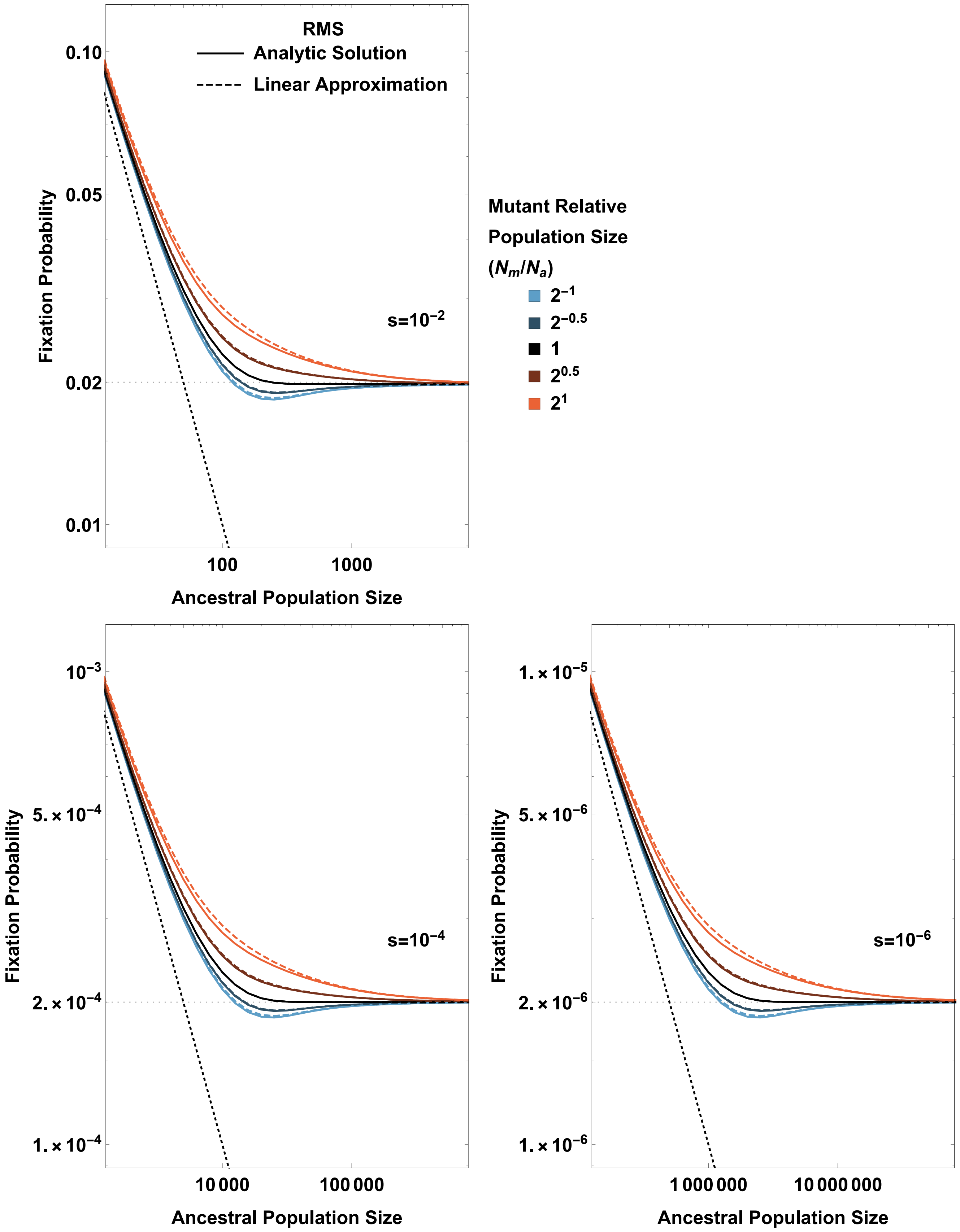


Figure S2: Fixation probability as a function of ancestral population size ($N_a$) for different values of the selection coefficients $s$ and relative boost (or drop) in population size ($N_m/N_a$), for the RMS model. The solid lines represent the analytical results, while the dashed lines represent the linear approximation results; both are closely matched with each other. The gray dotted line represents $\phi = 2s$ (the asymptotic limit of the fixation probability under positive selection at large population sizes), and the black dotted line represents $\phi = 1/N$ (the fixation probability under neutrality).

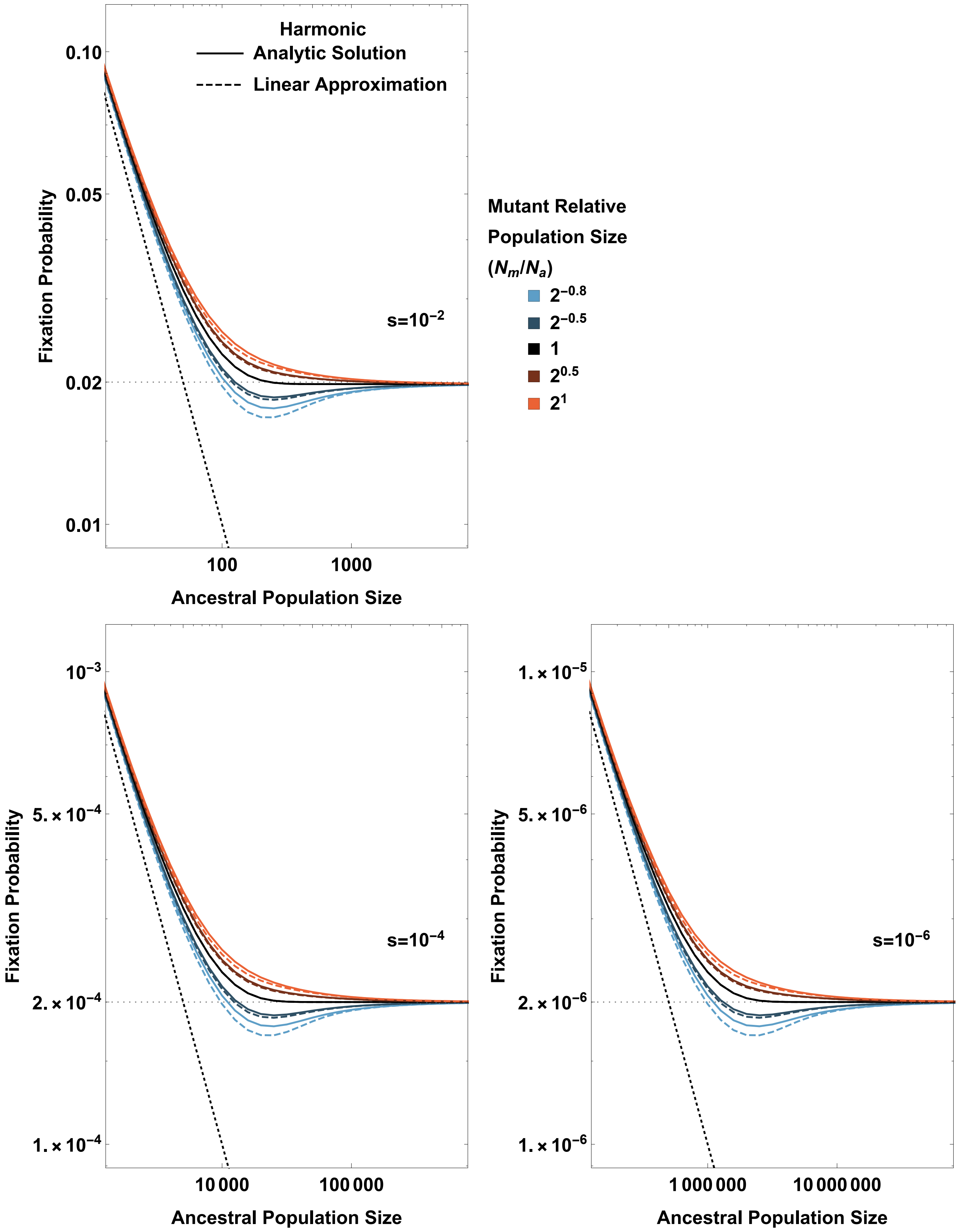


Figure S3: Fixation probability as a function of ancestral population size ($N_a$) for different values of the selection coefficients $s$ and relative boost (or drop) in population size ($N_m/N_a$), for the harmonic mean model. The solid lines represent the analytical results, while the dashed lines represent the linear approximation results. The gray dotted line represents $\phi = 2s$ (the asymptotic limit of the fixation probability under positive selection at large population sizes), and the black dotted line represents $\phi = 1/N$ (the fixation probability under neutrality).

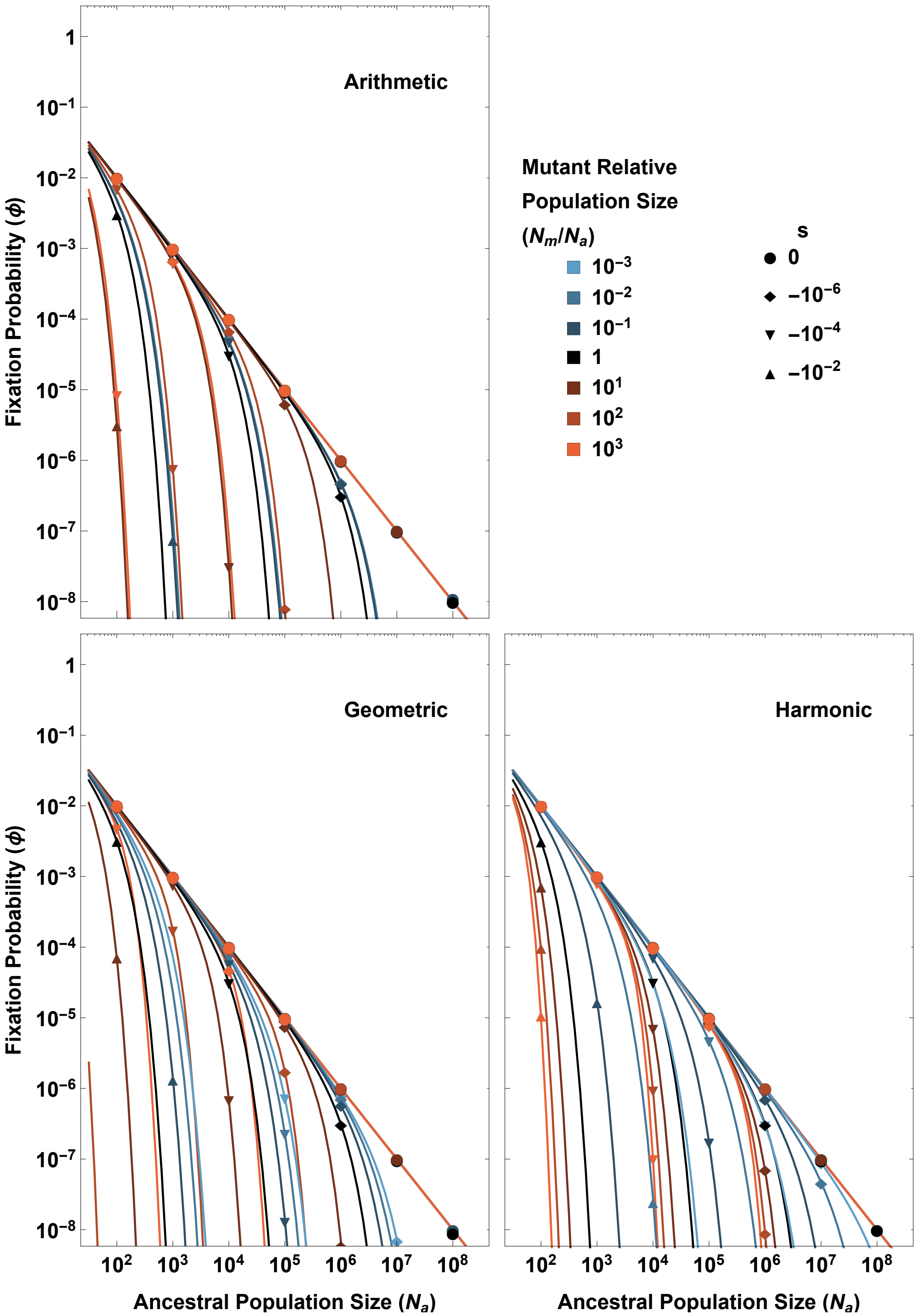


Figure S4: Fixation probability as a function of ancestral population size ($N_a$) for different values of negative selection coefficients $s$ (with different shapes) and relative boost (or drop) in population size ($N_m/N_a$, different colors), for different models. The solid lines represent the analytic results, while the points represent the simulation results. Note that our simulation results for negative $s$ had a strict lower bound on $N_m$ equal to 100.

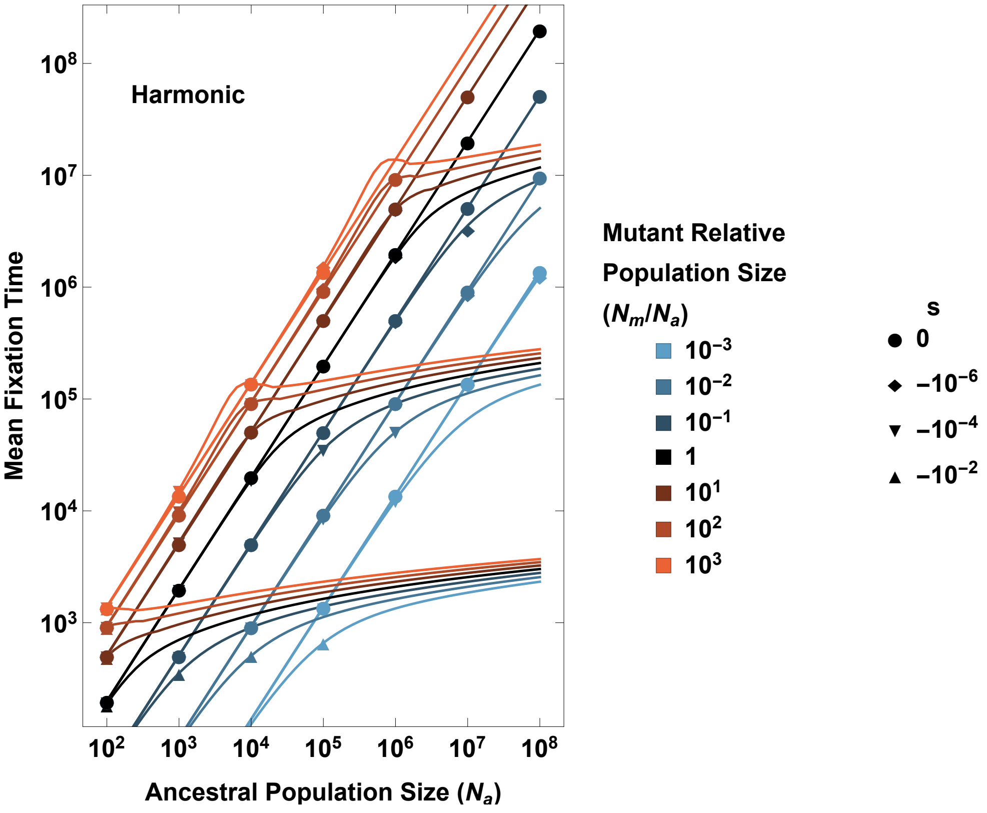


Figure S5: Mean fixation time as a function of ancestral population size ($N_a$) for different values of negative selection coefficients $s$ (with different shapes) and relative boost (or drop) in population size ($N_m/N_a$, different colors), for the harmonic mean model. The solid lines represent the analytic results, while the points represent the simulation results. Note that our simulation results for negative $s$ had a strict lower bound on $N_m$ equal to 100. Also, as the magnitude of $s$ increases and $N$ increases, the fixation probability significantly decreases. We have only plotted the simulation results for parameter values at which there were at least 100 runs that went to fixation out of a total of $5 \times 10^{10}$ runs.

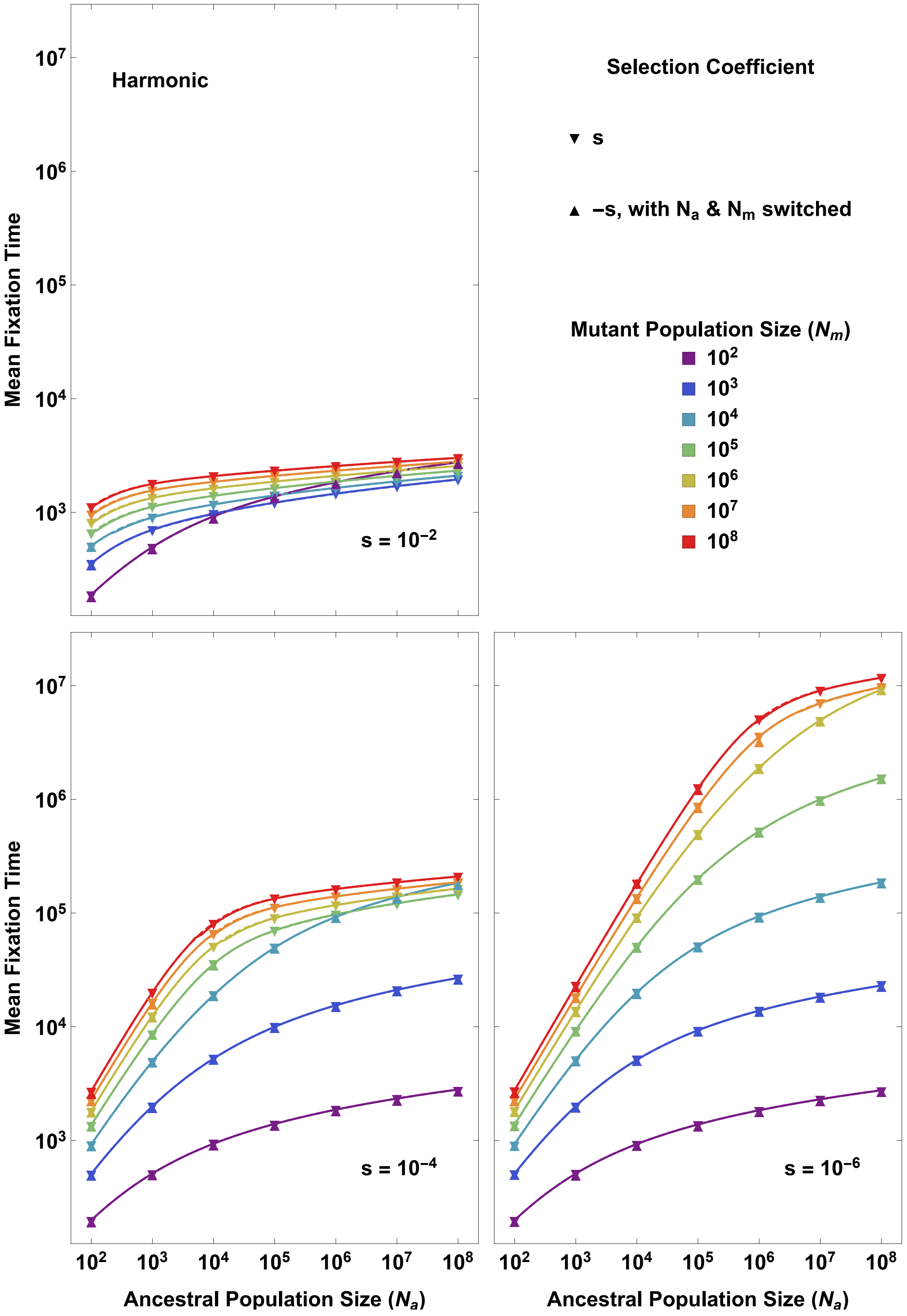


Figure S6: Mean fixation time (for harmonic mean model) as a function of ancestral population size ($N_a$) for different mutant population sizes ($N_m$) and different selection coefficients $s$. The downward triangle (▼) represents simulation with positive $s$ and the upward triangle (▲) represents simulation with negative $s$, with $N_a$ and $N_m$ switched. The hourglass shape appears as a consequence of a precise overlap between ▼ and ▲. Here, solid lines are the analytical results. As the magnitude of $s$ increases and $N$ increases, the fixation probability significantly decreases. We have only plotted the simulation results for parameter values at which there were at least 100 runs that went to fixation out of a total of $5 \times 10^{10}$ runs.

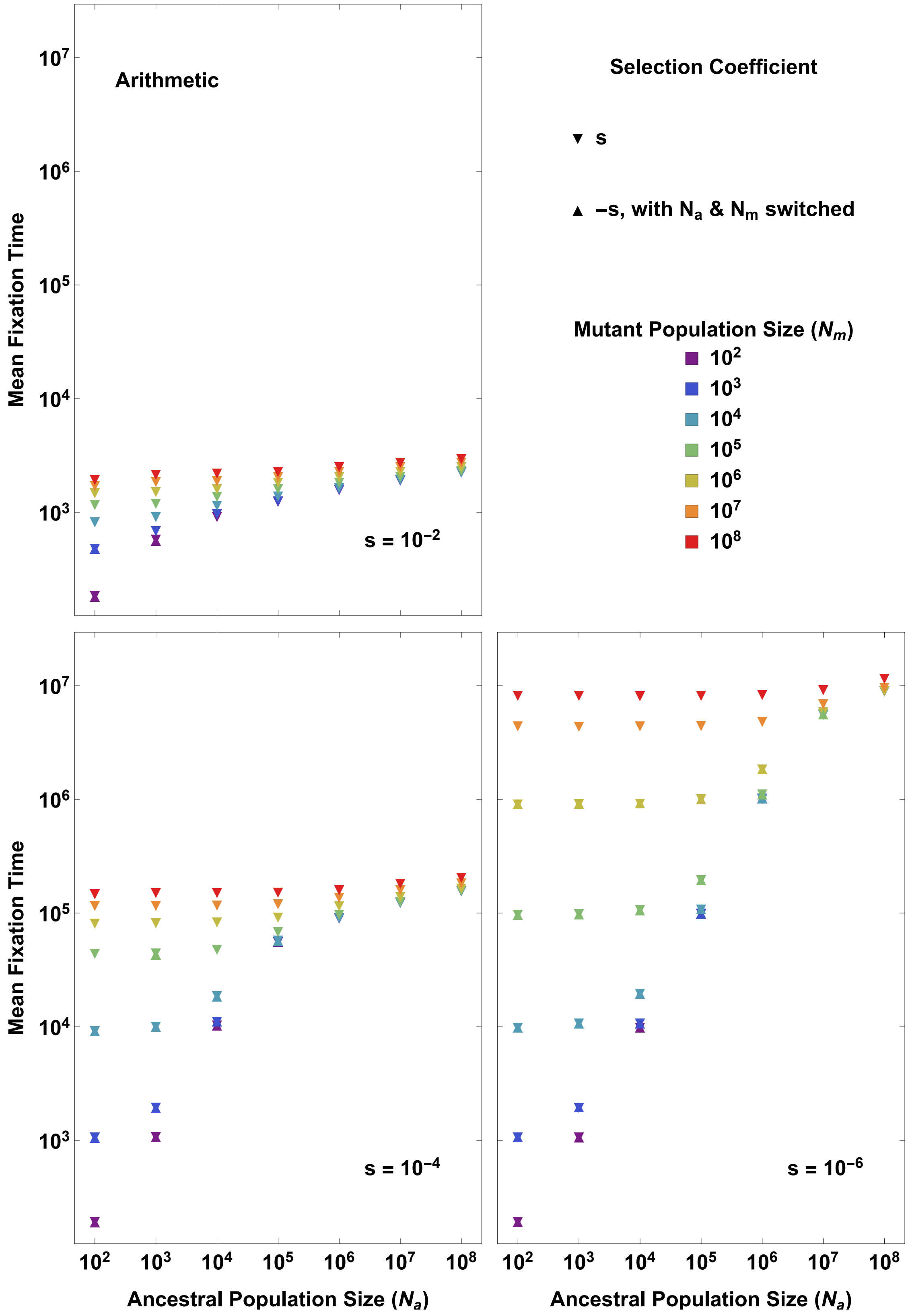


Figure S7: Mean fixation time (for arithmetic mean model) as a function of ancestral population size ($N_a$) for different mutant population sizes ($N_m$) and different selection coefficients $s$. The downward triangle (▼) represents simulation with positive $s$ and the upward triangle (▲) represents simulation with negative $s$, with $N_a$ and $N_m$ switched. The hourglass shape appears as a consequence of a precise overlap between ▼ and ▲. As the magnitude of $s$ increases and $N$ increases, the fixation probability significantly decreases. We have only plotted the simulation results for parameter values at which there were at least 100 runs that went to fixation out of a total of $5 \times 10^{10}$ runs.

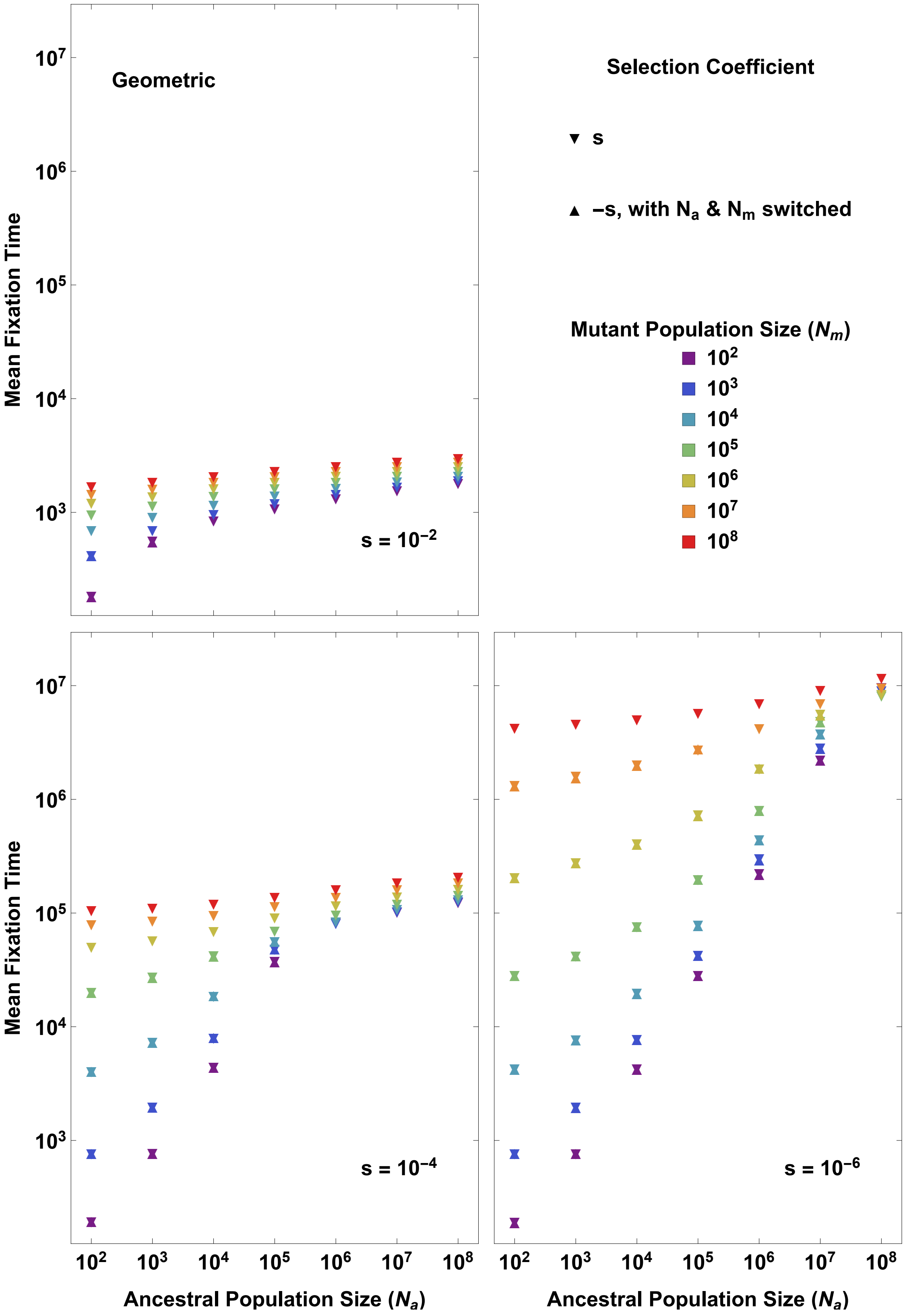


Figure S8: Mean fixation time (for geometric mean model) as a function of ancestral population size ($N_a$) for different mutant population sizes ($N_m$) and different selection coefficients $s$. The downward triangle (▼) represents simulation with positive $s$ and the upward triangle (▲) represents simulation with negative $s$, with $N_a$ and $N_m$ switched. The hourglass shape appears as a consequence of a precise overlap between ▼ and ▲. As the magnitude of $s$ increases and $N$ increases, the fixation probability significantly decreases. We have only plotted the simulation results for parameter values at which there were at least 100 runs that went to fixation out of a total of $5 \times 10^{10}$ runs.

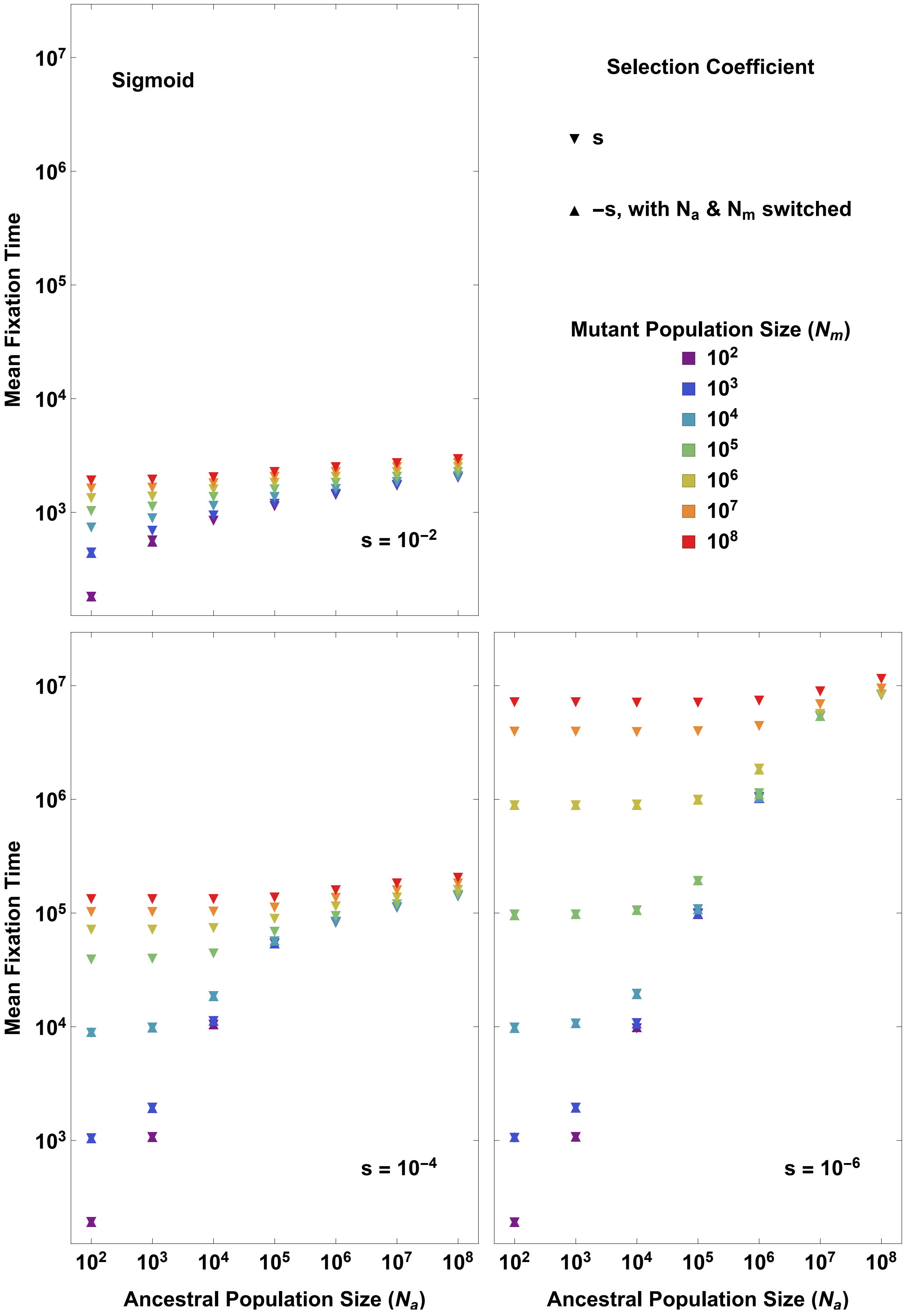


Figure S9: Mean fixation time (for sigmoid model) as a function of ancestral population size ($N_a$) for different mutant population sizes ($N_m$) and different selection coefficients $s$. The downward triangle (▼) represents simulation with positive $s$ and the upward triangle (▲) represents simulation with negative $s$, with $N_a$ and $N_m$ switched. The hourglass shape appears as a consequence of a precise overlap between ▼ and ▲. As the magnitude of $s$ increases and $N$ increases, the fixation probability significantly decreases. We have only plotted the simulation results for parameter values at which there were at least 100 runs that went to fixation out of a total of $5 \times 10^{10}$ runs.

#### Comparison to population growth as a function of time

In Kimura's logistic growth model [6], the population size grows deterministically with time as,

$$\frac{dN(t)}{dt} = rN\left(1 - \frac{N}{K}\right), \tag{S46}$$

where $K$ is the carrying capacity, and $r$ is the intrinsic growth rate of the population. The fixation probability for this model is given by [6],

$$\phi_K \approx 1 - e^{-2s\frac{s+r}{s+rN_a/K}} \approx 2s\frac{s+r}{s+rN_a/K}. \tag{S47}$$

We compare this to our linear model,

$$N(p_m) = N_a + N'(0)p_m,$$

with the same starting population size ($N_a$) and the same initial increase in population size as Kimura's model. In Kimura's model, the increase in population size in the first generation is given by,

$$\Delta N = rN_a\left(1 - \frac{N_a}{K}\right),$$

and as the first mutant allele arises, the mutant allele frequency changes from 0 to $1/N_a$, thus, in the first generation, $\Delta p_m = 1/N_a$. Thus, in our linear model, we set,

$$N'(0) = \frac{\Delta N}{\Delta p_m} = rN_a^2\left(1 - \frac{N_a}{K}\right). \tag{S48}$$

In the deterministic limit, $p_m$ grows as,

$$\frac{dp_m}{dt} = sp_m(1 - p_m), \tag{S49}$$

thus, the population size would grow as,

$$\frac{dN(t)}{dt} = s(N - N_a)\left(\frac{N_a + N'(0) - N}{N'(0)}\right). \tag{S50}$$

Since the rate of change of population size tends to 0 as $N$ approaches $N_a$, the above equation is not integrable with the initial condition $p_m = 0$ (corresponding to $N = N_a$) at $t = 0$. So, we instead integrate with the initial condition $p_m = 1/N_a$ at $t = 1$. This results in,

$$N(t) = N_a + \frac{N'(0)}{1 + (N_a - 1)e^{-\frac{s}{N'(0)}(t-1)}}. \tag{S51}$$

We note that there is no way to match the exact deterministic trajectory of Kimura's logistic growth model with our model, because comparing Eq. (S46) with Eq. (S50) shows that would require either $N_a = 0$ or $N'(0) = -N_a$ (implying $N(1) = 0$), both of which are not possible. In general, matching Kimura's model to the deterministic limit of any arbitrary model with mutant-allele dependent population size will not be possible, since combining Eq. (S46) and (S49) gives us,

$$\frac{dN(p_m)}{dp_m} = \frac{rN(1 - N/K)}{sp_m(1 - p_m)},$$

which is not integrable for the initial condition $N(p_m = 0) = N_a$. Specifically, the rate of change of $N$ tends to 0 as $p_m$ approaches 0 or 1 in models with frequency dependent population size, while the rate of change of $N$ maintains a finite non-zero value in Kimura's model. So, we instead compare our model with Kimura's by using the same starting population size ($N_a$) and the same initial increase in population size as Kimura's model. As a sanity check, plugging Eq. (S48) into Eq. (S51) yields,

$$N(t = 1) = N_a + rN_a\left(1 - \frac{N_a}{K}\right),$$

which approximately matches the expected population size after one generation in Kimura's model, given by Eq. (S46) with the initial condition $N(t = 0) = N_a$.

The analytic solution of the fixation probability for our linear model can be obtained from Table 1 in the main text, by replacing $N_m$ by $N_a + N'(0)$ in the row for the arithmetic model. We have compared the fixation probabilities from Kimura's model to our linear model for different parameter values in Fig. S10.

Now, from the last row of Table 1 in main text, we can obtain the following approximation for the fixation probability using the transformation $N_m \to N_a + N'(0)$, which is valid under the conditions listed in Table 1.

$$\phi \approx \frac{2}{N_a}\sqrt{\frac{sN'(0)}{\pi}} = 2\sqrt{\frac{sr}{\pi}\left(1 - \frac{N_a}{K}\right)}. \tag{S52}$$

This approximation is compared to the full analytic solution in Fig. S11, which shows that it holds good as long as $\phi \gg 2s$ in the regime of effective selection. Thus, the ratio of the fixation probabilities of the two models can be approximated as,

$$\frac{\phi}{\phi_K} \approx \frac{1}{\sqrt{c\pi}}\left(\frac{N_a/K + c}{1+c}\right)\sqrt{1 - \frac{N_a}{K}} \equiv f(c),$$

where $c = s/r$. Now, taking log and differentiating,

$$\frac{f'(c)}{f(c)} = -\frac{1}{2c} + \frac{1}{N_a/K + c} - \frac{1}{1+c}.$$

When $c \geq 1$, this is always negative. When $c < 1$, this is negative if

$$\frac{N_a}{K} > \frac{c(1-c)}{1+3c},$$

and positive if $N_a/K$ is less than this value. The results in the main text follow.

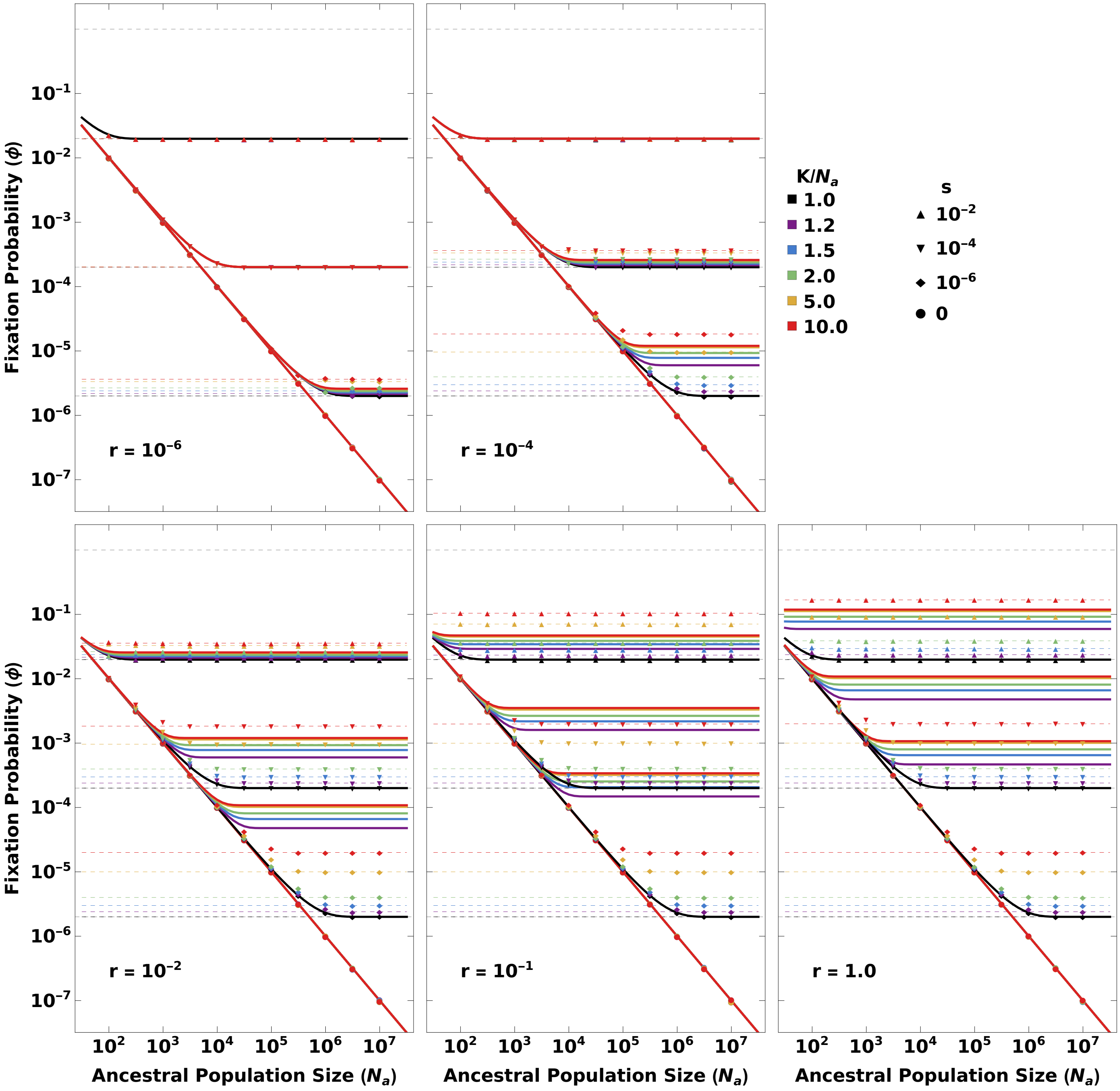


Figure S10: **A comparison between the fixation probabilities for Kimura's logistic growth model and our linear model.** Fixation probability is plotted as a function of ancestral population size ($N_a$) for different model parameters ($K$ and $r$), and different selection coefficients $s$ (represented by different markers). The points show the simulation results for Kimura's model, the dashed lines show Kimura's analytic solution for his model, and the solid lines show the analytic solution to our linear model with the same starting population size ($N_a$) and the same initial increase in population size as Kimura's model.

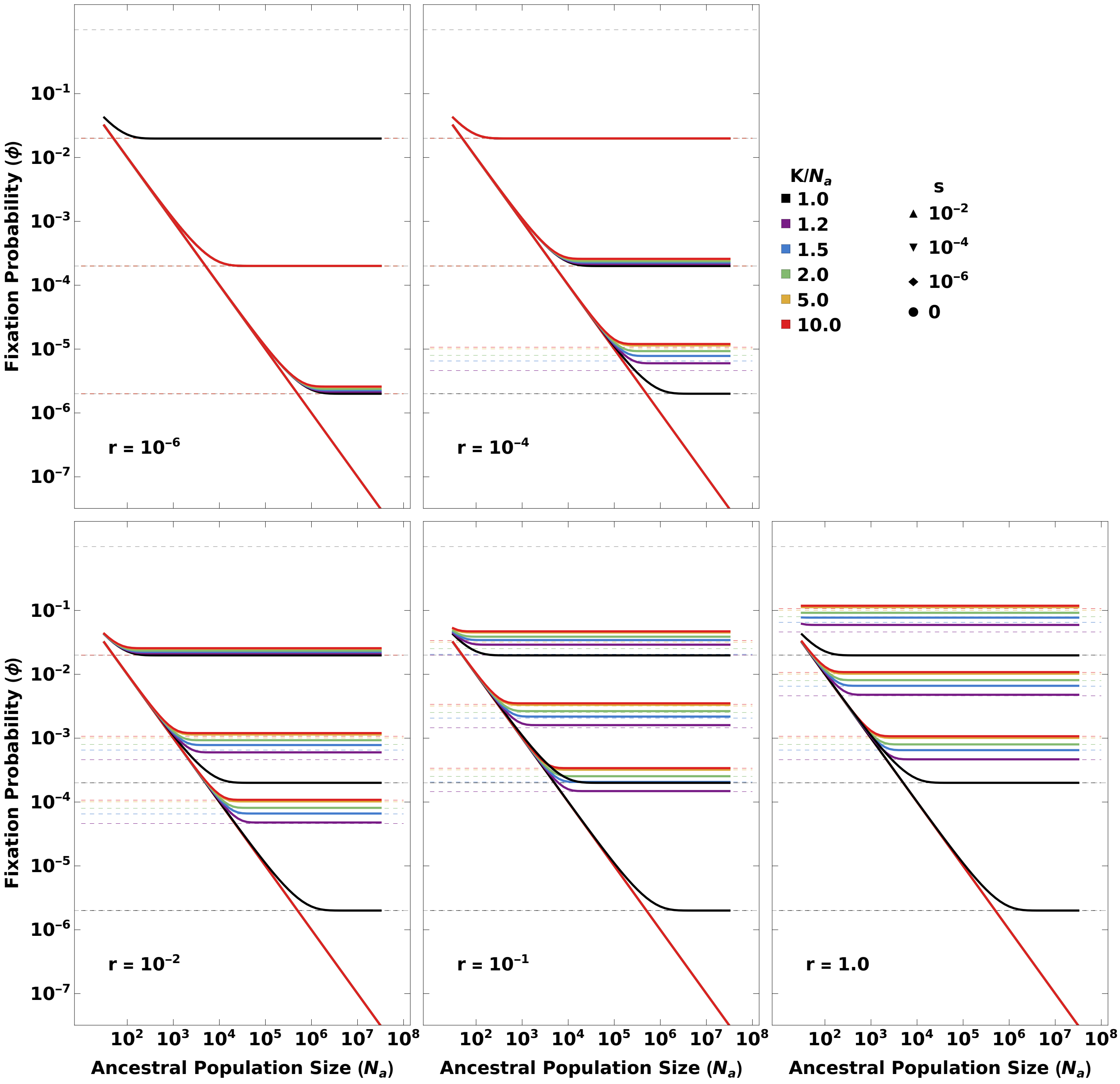


Figure S11: **Comparing the approximation to the full solution for the probability of fixation for our linear model.** Fixation probability is plotted as a function of ancestral population size ($N_a$) for different model parameters ($K$ and $r$), and different selection coefficients $s$ (represented by different markers). The solid lines show the analytic solution to our linear model with the same starting population size ($N_a$) and the same initial increase in population size as Kimura's model, same as in Fig. S10. The dashed lines show the maximum of the approximation given by Eq. S52 and $2s$. The approximation seems to work well as long as the fixation probability is $\gg 2s$ in the regime of effective selection.
